# CRISPR-based shuttle cloning of 1,397 human genes into UAS vectors

**DOI:** 10.1101/2025.02.16.638485

**Authors:** Xuelian Liu, Hanqing Xi, Miao Dai, Xiaoxue Li, Wen Xue, Guang Chen, Jialong Yan, Si Xu, Guifang Ou, Si Luo, Yonghong Tang, Ping Wei, Jiwu Wang

**Affiliations:** Clinical Research Institute, The Affiliated Nanhua Hospital, Hengyang Medical School, University of South China, Hengyang, Hunan 421002, China; Department of Neurology, The Affiliated Nanhua Hospital, Hengyang Medical School, University of South China, Hengyang, Hunan 421002, China; Institute of Biochemistry and Molecular Biology, Hengyang Medical School, University of South China, Hengyang, Hunan 421002, China; Shanghai Diabetes Institute, Shanghai Key Laboratory of Diabetes Mellitus, Shanghai Clinical Center for Diabetes, Shanghai Sixth People’s Hospital Affiliated to Shanghai Jiao Tong University School of Medicine, Shanghai 200233, China

**Keywords:** CRISPR/Cas9, DNA library, *Drosophila*, functional genomics, high-throughput cloning

## Abstract

The construction of genome-wide DNA libraries from publicly available resources is essential for leveraging functional genomics to investigate complex biological systems. However, all existing high-throughput cloning methods for transferring DNA fragments between vectors require PCR amplification of the DNA fragments, rendering the construction of genome-wide DNA libraries labor-intensive and time-consuming. By introducing a concept of CRISPRshuttle cassette, we herein present a method named CRISPR-based shuttle cloning (CRISPRshuttle cloning). This method enables the high-throughput transfer of numerous DNA fragments from original plasmids with identical backbones to a different vector background without the need for PCR amplification of the DNA fragments. The procedure comprises two-step test tube reactions followed by bacterial transformation. Using CRISPRshuttle we successfully generated a library of GAL4/UAS-based UAS-ORF plasmids covering 1,397 human genes conserved in *Drosophila*. This library may serve as a valuable resource for gain-of-function screening in cultured cells and for the creation of a transgenic UAS-ORF library in *Drosophila*.

---

Functional genomics is a powerful tool for elucidating the function of all genes in an organism and primarily relies on construction and manipulation of genome-wide DNA libraries such as cDNAs, ORFs, gene promoters and inverted repeats for RNAi. Typically, a DNA library must be transferred to a destination vector in a high-throughput manner for functional genomics studies. However, the construction of genome-wide DNA libraries, such as cDNA/ORF overexpression libraries, has been challenging due to limitations in high-throughput DNA cloning methods. This severely restricts the utility of functional genomics. The prevailing high-throughput cloning methods for constructing and manipulating genome-wide DNA libraries primarily include Gateway, In-Fusion, Creator, and Univector cloning systems, all of which are based on site-specific recombination. Among these, the Gateway cloning system is the most extensively employed high-throughput cloning method.

All the aforementioned high-throughput cloning methods predominantly rely on PCR amplification of the DNA fragments of interest. This step requires individualized manipulations for each DNA fragment, including primer design and synthesis, gel purification, and DNA sequencing, which are laborious and time-consuming. Additionally, PCR amplification is particularly problematic for long DNA fragments. Consequently, the PCR amplification of DNA fragments of interest is not only costly but also a rate-limiting step in high-throughput cloning. For example, although cDNA and ORF resources for human, mouse and *Drosophila* have been publicly available for nearly two decades, the construction of a genome-wide GAL4/UAS (upstream activating sequence)-based UAS-cDNA/ORF plasmid library from these resources has been severely impeded by the PCR amplification of cDNAs and ORFs[1].

We previously developed a high-throughput cloning method, CRISPRmass, for constructing a genome-wide UAS-cDNA/ORF plasmid library from publicly available cDNA/ORF resources[2]. However, CRISPRmass is applicable solely to the insertion of an identical DNA fragment (e.g., a UAS module) into the identical backbones of different plasmids[2]. It does not allow for the transfer of DNA fragments (e.g., cDNAs or ORFs) between vectors, thereby limiting its utility in DNA cloning.

By introducing the concept of a CRISPRshuttle cassette, we developed a novel high-throughput DNA cloning method termed CRISPR-based shuttle cloning (CRISPRshuttle cloning). This method allows for the transfer of numerous DNA fragments of interest from original plasmids with identical backbones to a different vector background through two-step test tube reactions prior to bacterial transformation, thereby eliminating the need for PCR amplification of the DNA fragments (Fig. 1A). In the first-step test tube reaction, different DNA fragments of interest are excised from their original plasmids by digesting the plasmid backbones with Cas9/sgRNA 1 and Cas9/sgRNA 2. Cas9/sgRNA 1 targets the backbone sequence adjacent to the 5’ end of the DNA fragments, while Cas9/sgRNA 2 targets the backbone sequence adjacent to the 3’ end. The released DNA fragments do not need to be purified, and the reaction products can be directly used in the second-step test tube reaction. In the second-step test tube reaction, the released DNA fragments are transferred to the CRISPRshuttle cassette of a CRISPRshuttle-compatible destination vector via Gibson assembly, yielding the desired plasmids. A CRISPRshuttle cassette consists of approximately 20-40 bp of backbone sequence flanking both the 5’ and 3’ ends of the DNA fragments, and one or two unique restriction enzyme recognition sites between these sequences. These recognition sites are used for linearizing the CRISPRshuttle-compatible destination vector and must be unique within the vector. The CRISPRshuttle-compatible destination vector is created by cloning the CRISPRshuttle cassette into a destination vector, which is then linearized by the restriction enzymes 1 and 2 between the two 20-40 bp backbone sequences of the CRISPRshuttle cassette prior to the second-step test tube reaction. The antibiotic resistance gene of the destination vector must differ from that of the original plasmids; if they are identical, it has to be replaced with a distinct one. Products of the second-step test tube reaction are directly subjected to bacterial transformation without purification. Bacterial transformants are selected on LB plates containing the antibiotic corresponding to the antibiotic resistance gene of the destination vector. Theoretically only the colonies harboring the desired plasmids can grow on the LB plates.

**Figure 1.**
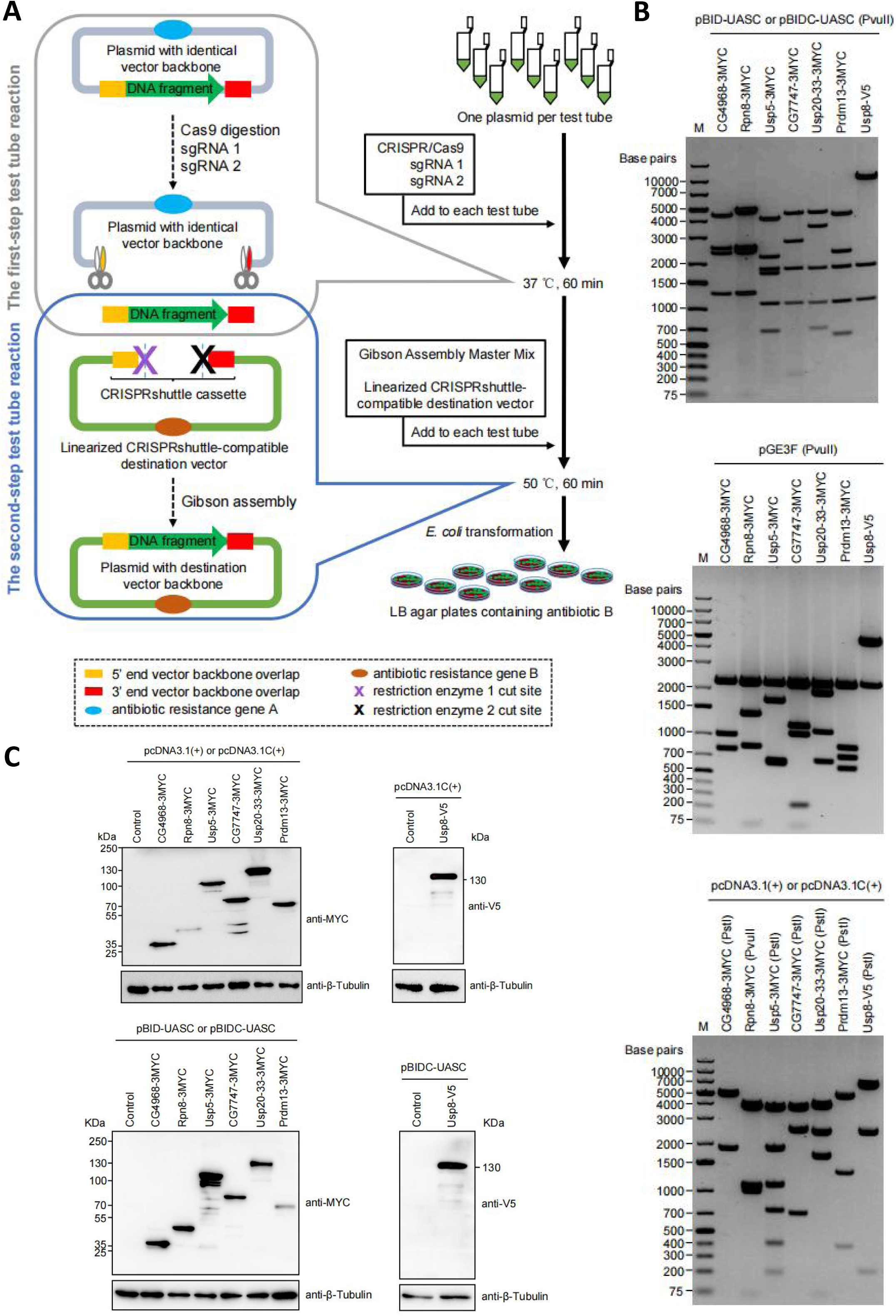
CRISPRshuttle enables high-throughput transfer of DNA fragments between vectors. **(A)** The CRISPRshuttle pipeline. The first-step test tube reaction is to cleave the identical vector backbone of the original plasmids using Cas9 in conjunction with sgRNA 1 and sgRNA 2. This step is to release the DNA fragments of interest from the vector backbone. The Cas9 cleavage sites are located on the backbone sequences adjacent to the 5’ end and 3’ end of the DNA fragments of interest respectively. The released DNA fragments do not require purification and can be directly used for the second-step test tube reaction. The second-step test tube reaction is to clone the released DNA fragments into a linearized CRISPRshuttle-compatible destination vector using Gibson assembly, yielding the desired plasmids. The Gibson assembly products, without purification, are directly subjected to *E. coli* transformation. Transformants are selected on LB plates containing an antibiotic corresponding to the antibiotic resistance gene (brown ellipse) of the destination vector. **(B)** Restriction analysis of 21 constructs generated by CRISPRshuttle cloning. The seven cDNAs and ORFs cloned in seven original vectors were transferred to three destination vectors using CRISPRshuttle respectively, resulting in the 21 constructs. The expected PvuII digestion patterns of all the 21 constructs were observed. (M) DNA marker. (**C**) Western blotting showed the seven cDNAs and ORFs of the 21 constructs generated using CRISPRshuttle were expressed in HEK293 cells.

To assess the feasibility of utilizing CRISPRshuttle for DNA fragment transfer between vectors, we attempted to transfer seven cDNAs and ORFs from original plasmids with distinct backbones to three destination vectors respectively using CRISPRshuttle, thereby generating a total of 21 constructs. The vectors of these seven original plasmids include six vectors from the *Drosophila* Genomics Resource Center (DGRC) Gold Collection, namely pOT2, pOTB7, pFLC-I, pBS SK(−), pCR2.1, pDNR-Dual, and one vector from the human CCSB-Broad Lentiviral Expression Library, pLX304. The proteins encoded by the seven cDNAs and ORFs in the original plasmids are tagged with a 3xMYC or V5 epitope at their C termini[2]. The three destination vectors include a GAL4/UAS-based UAS vector, pBID-UASC[3], a mammalian expression vector, pcDNA3.1(+), and a Gateway entry vector, pGE3F[4].

To release the seven cDNAs and ORFs from their original plasmids, we first identified the optimal Cas9 cleavage sites on the plasmid backbones. We evaluated the in vitro cleavage efficiency of all candidate sgRNAs (Supplementary Table S1; Supplementary Fig. S1) and selected an optimal sgRNA for each original vector for subsequent experiments (Supplementary Table S2).

To prepare CRISPRshuttle cassettes for the seven original vectors, we synthesized seven pairs of PAGE-purified DNA oligonucleotides (Supplementary Table S3). These oligonucleotides were annealed to form seven CRISPRshuttle cassettes, which were subsequently ligated into the EcoRI and XbaI sites of the multiple cloning sites of the corresponding three destination vectors respectively. This process yielded 21 CRISPRshuttle-compatible destination vectors (Supplementary Table S4).

We next performed two-step test tube reactions to transfer the seven cDNAs and ORFs from their original plasmids to 21 CRISPRshuttle-compatible destination vectors using CRISPRshuttle. In the first-step test tube reactions, the seven cDNAs and ORFs were released from their original plasmids via Cas9/sgRNA digestion. In the second-step test tube reactions, the released cDNAs and ORFs, without purification, were directly ligated into the 21 EcoRI/XhoI-linearized CRISPRshuttle-compatible destination vectors using Gibson assembly. The Gibson assembly products were then directly transformed into *E. coli*. Restriction analysis confirmed that the 21 resulting plasmids were successfully constructed via CRISPRshuttle (Fig. 1B; Supplementary Table S4). Subsequent Western blotting analysis detected the expected molecular weights of the proteins encoded by the seven cDNAs and ORFs in 14 of the resulting plasmids in the transfected cells (Fig. 1C). Collectively, these results demonstrate that CRISPRshuttle can be utilized to transfer a DNA fragment of interest between vectors, eliminating the need for PCR amplification of the DNA fragment.

Next, we explored whether CRISPRshuttle could be applied for high-throughput cloning. Drosophila has proven to be a powerful genetic model system for studying genes involved in various biological processes. The majority of Drosophila genes are evolutionarily conserved between flies and humans. Numerous studies have elucidated the functions of human genes through creating transgenic flies expressing human genes. The DGRC Gold Collection represents the most comprehensive and publicly available Drosophila cDNA/ORF plasmid library covering approximately 10,000 protein-coding genes in Drosophila. We selected 1,766 ORF clones from the human CCSB-Broad Lentiviral Expression Library[5], which are conserved in Drosophila based on OrthoDB v9.1. These human genes were chosen because their orthologs in Drosophila are not included in the DGRC Gold Collection, and thus their functions had not been systematically characterized. Among the selected 1,766 ORFs clones, 178 clones either failed to grow in LB medium or lacked the pLX304-specific restriction digestion patterns. Utilizing the remaining 1,588 ORF clones as original plasmids, we applied CRISPRshuttle to construct a UAS-ORF plasmid library using two GAL4/UAS-based UAS vectors, pBIDC-UASC-pLXvect and p10UASTattB-Cam-pLXvect, as CRISPRshuttle-compatible destination vectors (Fig. 2A). Following *E. coli* transformation, at least one colony grew on chloramphenicol-containing LB plates for all 1,588 CRISPRshuttle reactions. We randomly picked one or two colonies from each plate for plasmid DNA extraction. Subsequent restriction digestions with PvuII confirmed the presence of the expected fragments (Fig. 2B, Supplementary Table S5).

**Figure 2.**
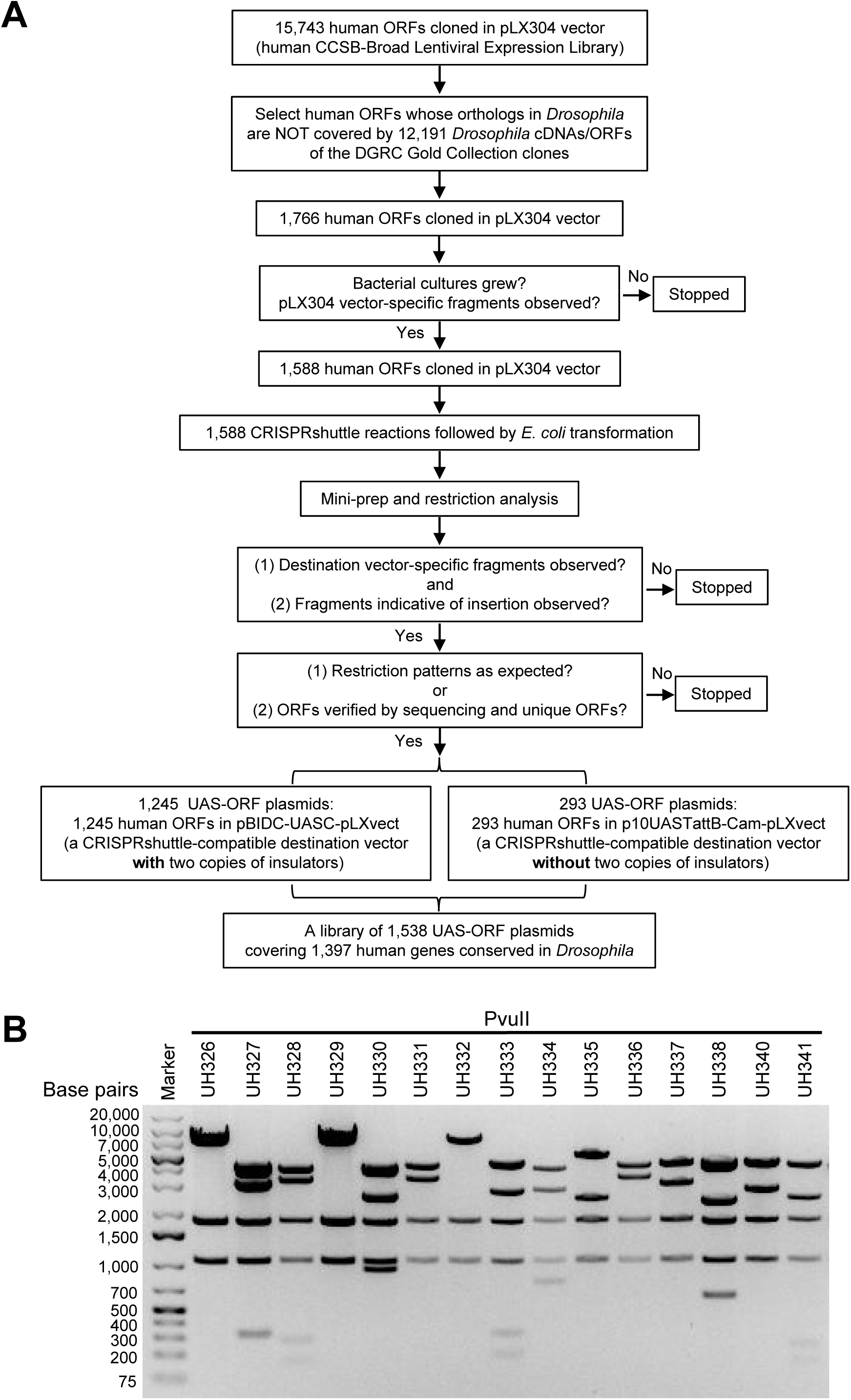
Generation of 1,538 UAS-ORF constructs covering 1,397 human genes conserved in *Drosophila* using CRISPRshuttle cloning. **(A)** 1,538 ORFs from the human CCSB-Broad Lentiviral Expression Library were transferred to two GAL4/UAS-based UAS vectors, pBIDC-UASC-pLXvect and p10UASTattB-Cam-pLXvect, using CRISPRshuttle. These two UAS vectors serve as CRISPRshuttle-compatible destination vectors. **(B)** Representative results of restriction analysis of UAS-ORF constructs that were generated by CRISPRshuttle-assisted transfer of human ORFs cloned in pLX304 to pBIDC-UASC-pLXvect. 15 UAS-ORF constructs were digested by PvuII and analyzed on 0.8% agarose gel, and the expected restriction patterns for these constructs were observed (Supplemental Table S5).

In summary, we successfully constructed a library of 1,538 UAS-ORF plasmids using CRISPRshuttle. Specifically, 1,538 human ORFs were transferred to the pBIDC-UASC-pLXvect vector, and 293 human ORFs were transferred to the p10UASTattB-Cam-pLXvect vector. The library comprises plasmids for 1,397 human genes, with 131 genes represented by 2-3 isoforms (Fig. 2A). This library can potentially serve as a valuable resource for gain-of-function screening in cultured cells and for the creation of a transgenic UAS-ORF library in Drosophila.

Furthermore, we evaluated the efficiency of CRISPRshuttle (Supplementary Fig. S2). The efficiency of CRISPRshuttle reaches 94.5% for the transfer of DNA fragments to CRISPRshuttle-compatible destination vectors bearing two repetitive sequences, and 96.1% for the transfer of DNA fragments to destination vectors lacking repetitive sequences. It is noteworthy that approximately 300 UAS-ORF plasmids were successfully generated using CRISPRshuttle cloning by two researchers within seven days, demonstrating the suitability of CRISPRshuttle for high-throughput cloning.

Finally, we compared CRISPRshuttle with CRISPRmass and Gateway (Table 1). Notably, CRISPRshuttle significantly outperformed Gateway in transferring DNA fragments between vectors.

**Table 1.** Comparison of CRISPRshuttle with CRISPRmass and Gateway via construction of a library of 1,538 UAS-ORF plasmids from 1,538 ORF clones of the human CCSB-Broad Lentiviral Expression Library.

|  |  | Gateway | CRISPRmass | CRISPRshuttle |
| --- | --- | --- | --- | --- |
| For generating entry constructs | Number of entry constructs generated | 1,538 | Not applicable | Not applicable |
|  | Pairs of primers designed for PCR amplification of every single ORF | Minimum 1,538 | Not applicable | Not applicable |
|  | PCR reactions for every single ORF | Minimum 1,538 | Not applicable | Not applicable |
|  | Samples for agarose gel electrophoresis for every single cDNA/ORF | Minimum 1,538 | Not applicable | Not applicable |
|  | Gel purifications for every single ORF | Minimum 1,538 | Not applicable | Not applicable |
|  | Efficiency of BP reaction | >90%<sup>1</sup> | Not applicable | Not applicable |
|  | Colonies to be analyzed for entry constructs | More than 1,538 | Not applicable | Not applicable |
|  | Sequencing reactions for every single ORF | More than 1,538 | Not applicable | Not applicable |
|  | ORF-specific primers for sequencing | Required for >2-kb ORF | Not applicable | Not applicable |
| For generating UAS-ORF constructs | sgRNAs used | Not applicable | 2 | 2 |
|  | Gateway-compatible destination vector to be constructed | 1 | Not applicable | Not applicable |
|  | UAS module to be constructed | Not applicable | 1 | Not applicable |
|  | CRISPRshuttle-compatible destination vector to be constructed | Not applicable | Not applicable | 1 |
|  | Reactions | LR reaction | CRISPR/Cas9 digestion, Gibson assembly | CRISPR/Cas9 digestion, Gibson assembly |
|  | Colonies to be analyzed for UAS-ORF constructs | 1,538 | 1,538 | 1,538 |
|  | Vector backbone changed | Yes | No | Yes |
|  | Efficiency | >95%<sup>2</sup> | 98.6%<sup>3</sup> | >94.5%<sup>4</sup> |

<sup>1</sup>Based on Gateway Recombination Cloning brochure (Thermo Fisher Scientific; https://www.thermofisher.com/cn/zh/home/life-science/cloning/gateway-cloning.html).
<sup>2</sup>This is the LR reaction efficiency based on Gateway Recombination Cloning brochure (Thermo Fisher Scientific; https://www.thermofisher.com/cn/zh/home/life-science/cloning/gateway-cloning.html).
<sup>3</sup>Based on the study in which UAS modules were inserted into 98.6% of 5,987cDNA/ORF plasmids successfully under the condition of randomly analyzing only one colony for each UAS-cDNA/ORF construct. (Wei et al. Genome Research, 2020)
<sup>4</sup>Based on the current study in which 94.5% of 697 UAS-ORF plasmids were successfully constructed by CRISPRshuttle under the condition of randomly analyzing only one colony for each UAS-ORF construct.

The CRISPR/Cas9 enzyme utilized in the CRISPRshuttle method can be substituted with other CRISPR/Cas enzymes, such as SpRY and Cas12a[6–8]. Our findings indicate that CRISPRshuttle is likely to be broadly applicable for constructing genome-wide DNA libraries and individual plasmids.

## Acknowledgements

We thank Xiaohui Cai for the technical assistance.

## Funding

This work was supported by the National Natural Science Foundation of China (32071135) and the Scientific Research Fund for Talented Scholars, The Affiliated Nanhua Hospital, Hengyang Medical School, University of South China.

## Conflict of Interest

A patent application has been filed for this study.

---

**Supplementary Figure S1.**
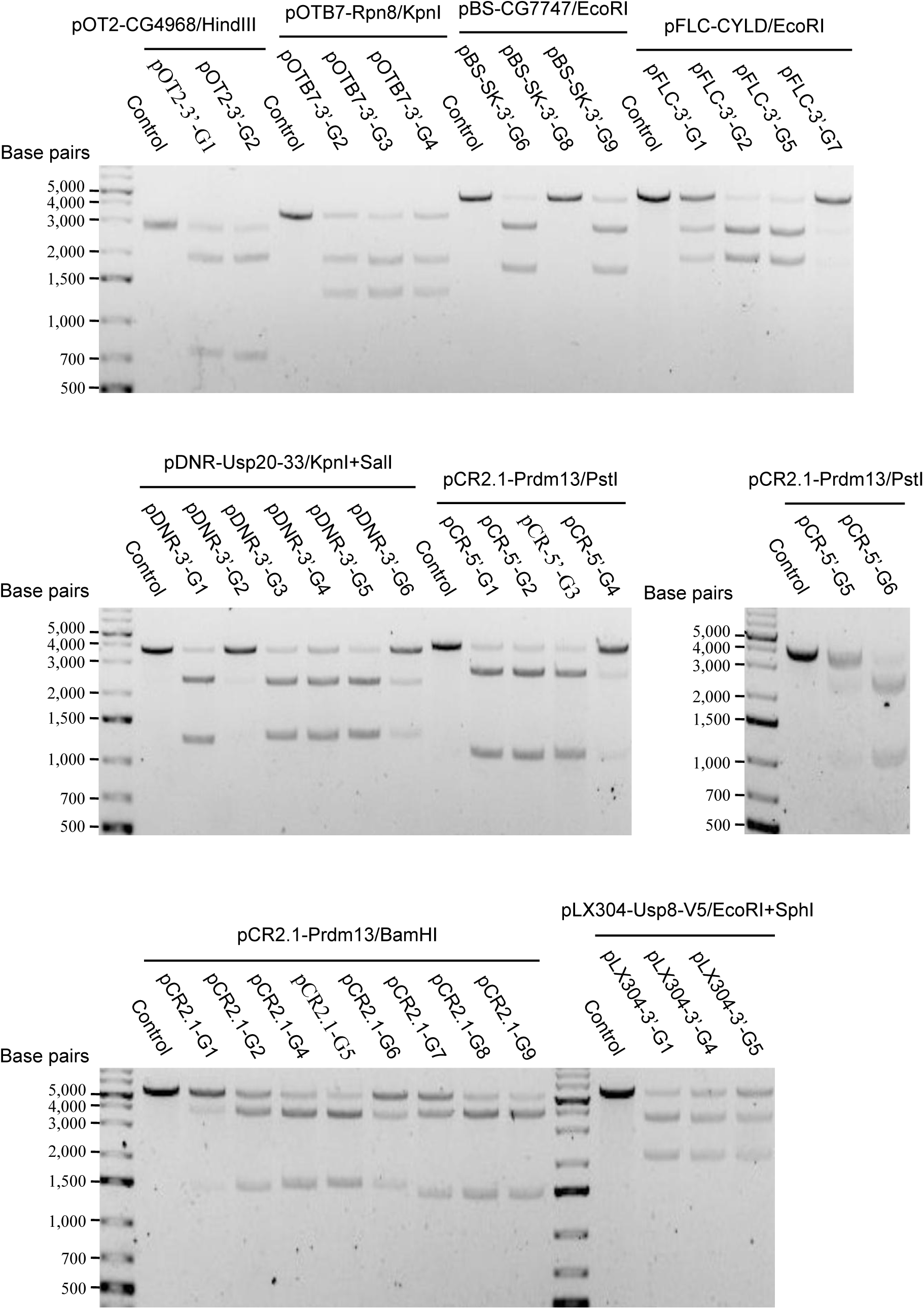
Evaluation of sgRNAs that target vector backbone sequences adjacent to the DNA fragments. These vectors include pOT2, pOTB7, pFLC-I, pBS SK(−), pCR2.1, and pDNR-Dual from the DGRC Gold Collection, and pLX304 from the human CCSB-Broad Lentiviral Expression Library. The sgRNAs and DNA substrates are listed in Supplementary Table S1.

**Supplementary Figure S2.**
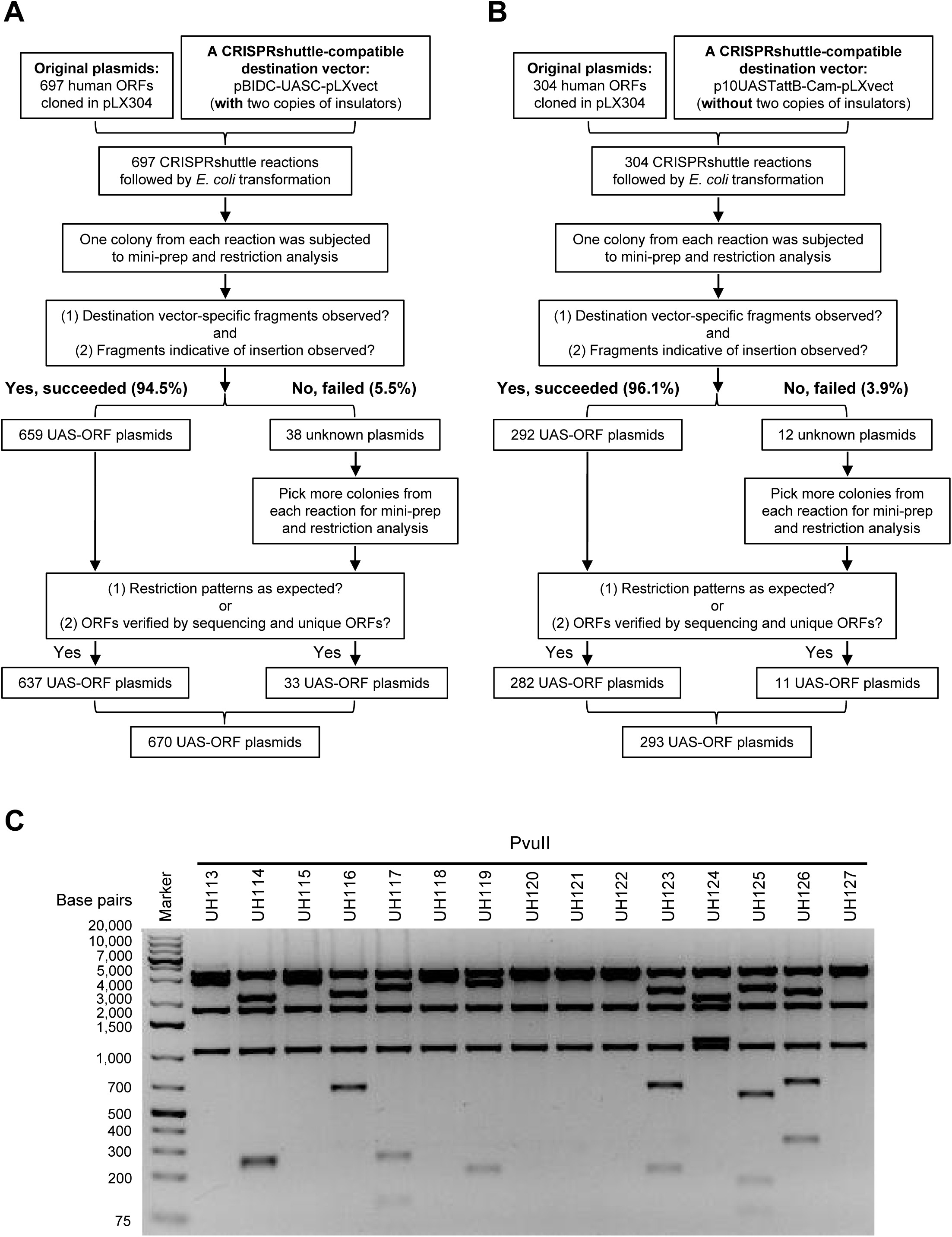
Evaluation of the efficiency of CRISPRshuttle cloning. (**A**) 697 CRISPRshuttle reactions were set up to transfer 697 human ORFs cloned in pLX304 vector to a CRISPRshuttle-compatible destination vector, pBIDC-UASC-pLXvect, which contains repetitive sequences. 659 human ORFs of the original plasmids were successfully transferred to pBIDC-UASC-pLXvect, resulting in 659 UAS-ORF plasmids. (**B**) 304 CRISPRshuttle reactions were set up to transfer 304 human ORFs cloned in pLX304 vector to a CRISPRshuttle-compatible destination vector, p10UASTattB-Cam-pLXvect, which does not contain repetitive sequences. 292 human ORFs of the original plasmids were successfully transferred to p10UASTattB-Cam-pLXvect, resulting in 292 UAS-ORF plasmids. (**C**) Representative results of restriction analysis of UAS-ORF constructs that were generated by CRISPRshuttle-assisted transfer of human ORFs cloned in pLX304 to p10UASTattB-Cam-pLXvect. 15 UAS-ORF constructs were digested by PvuII and analyzed on 0.8% agarose gel, and the expected restriction patterns for these constructs were observed (Supplemental Table S5).

**Supplementary Table S1.**
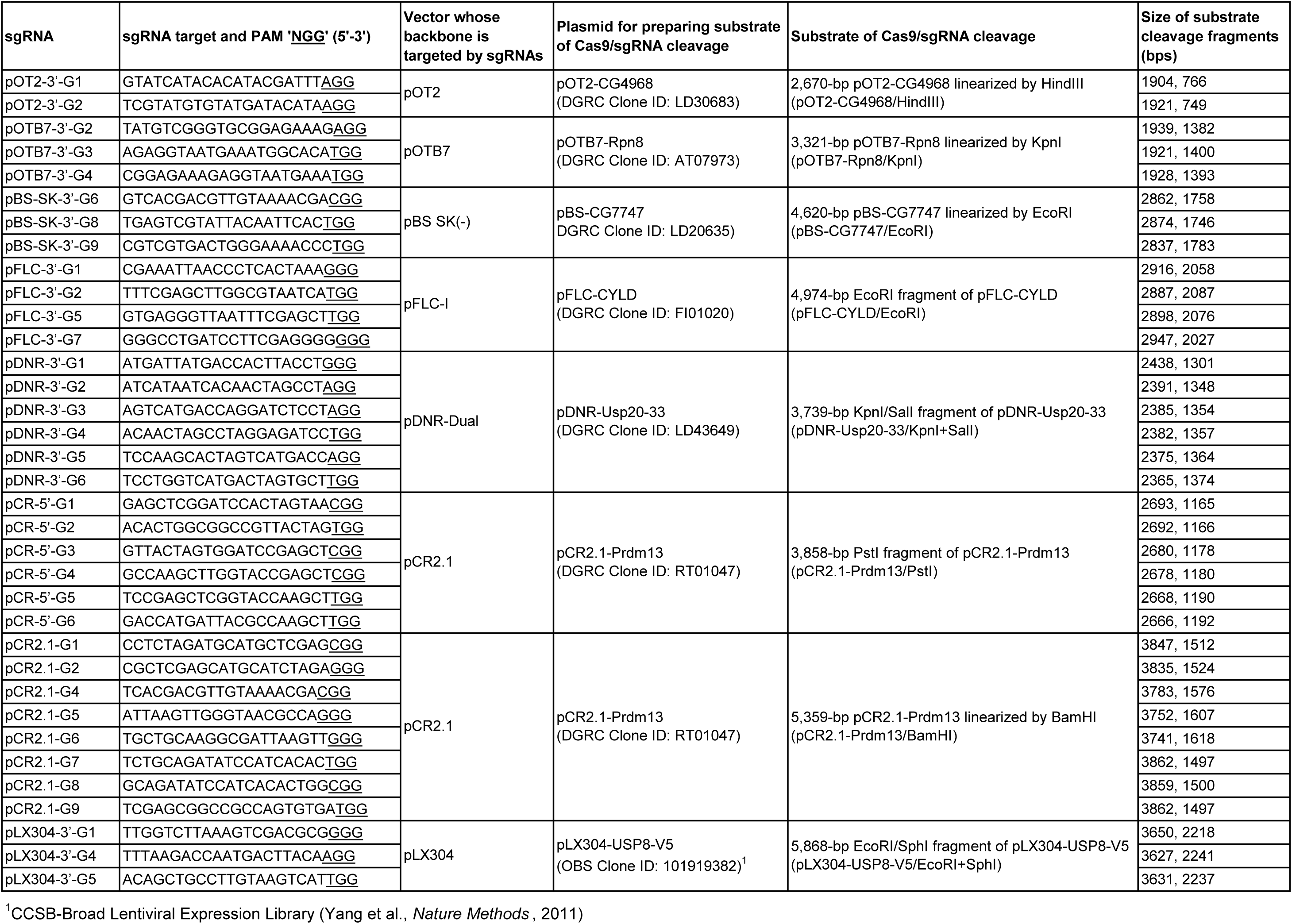
Information for sgRNA evaluation.

**Supplementary Table S2.**
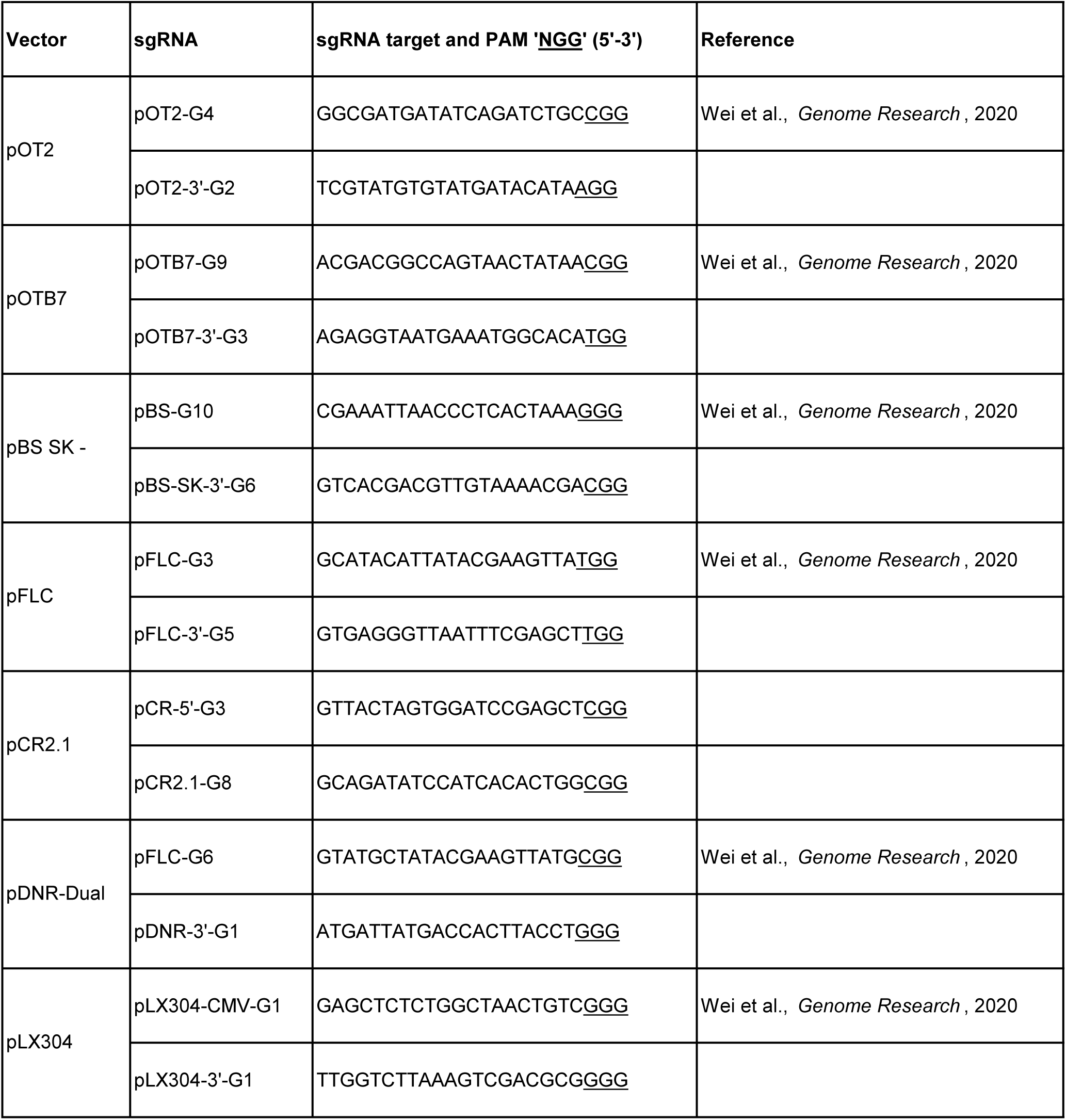
sgRNAs used for CRISPRshuttle.

**Supplementary Table S3.**
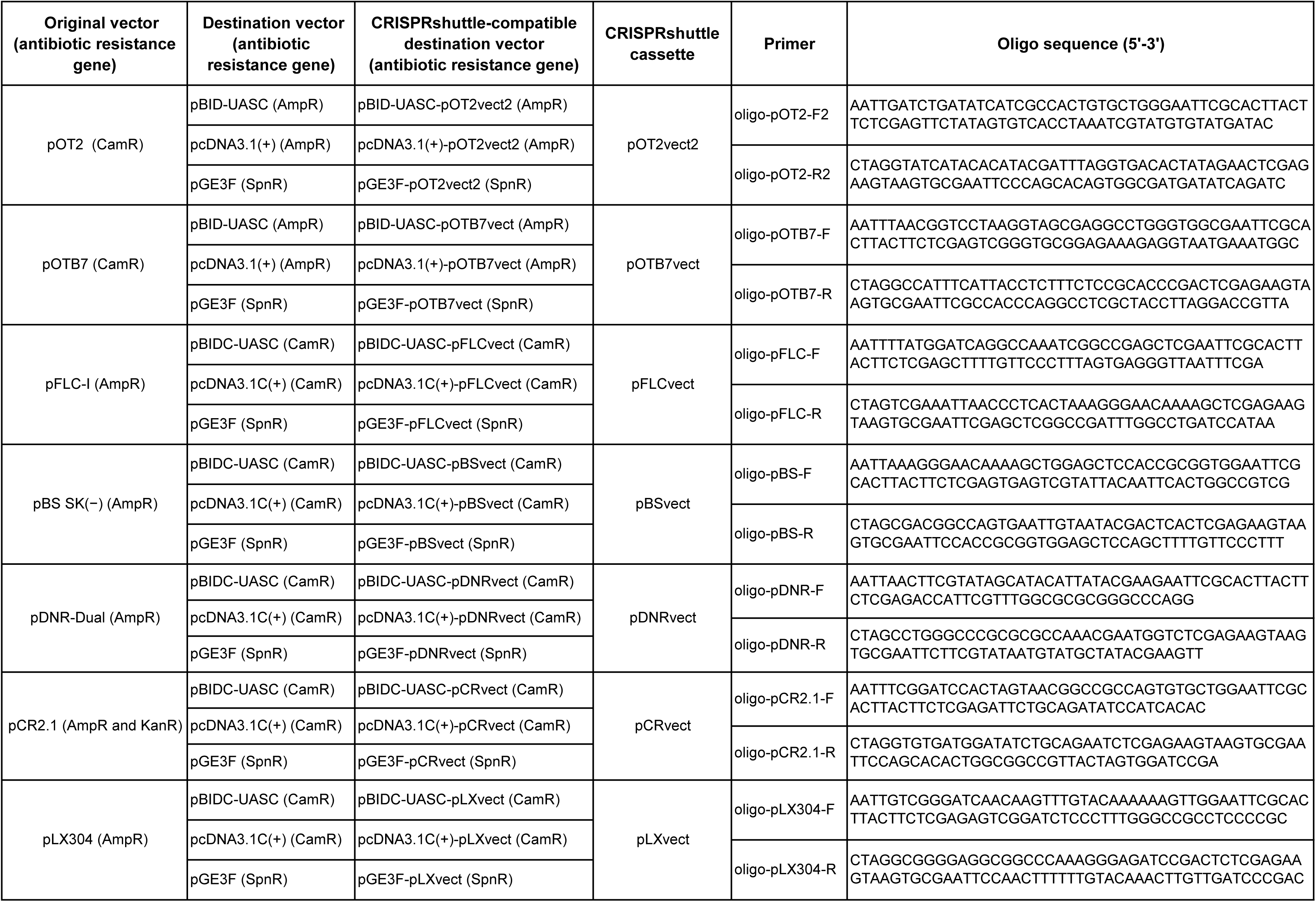
Oligos used for construction of CRISPRshuttle cassettes.

**Supplementary Table S4.**
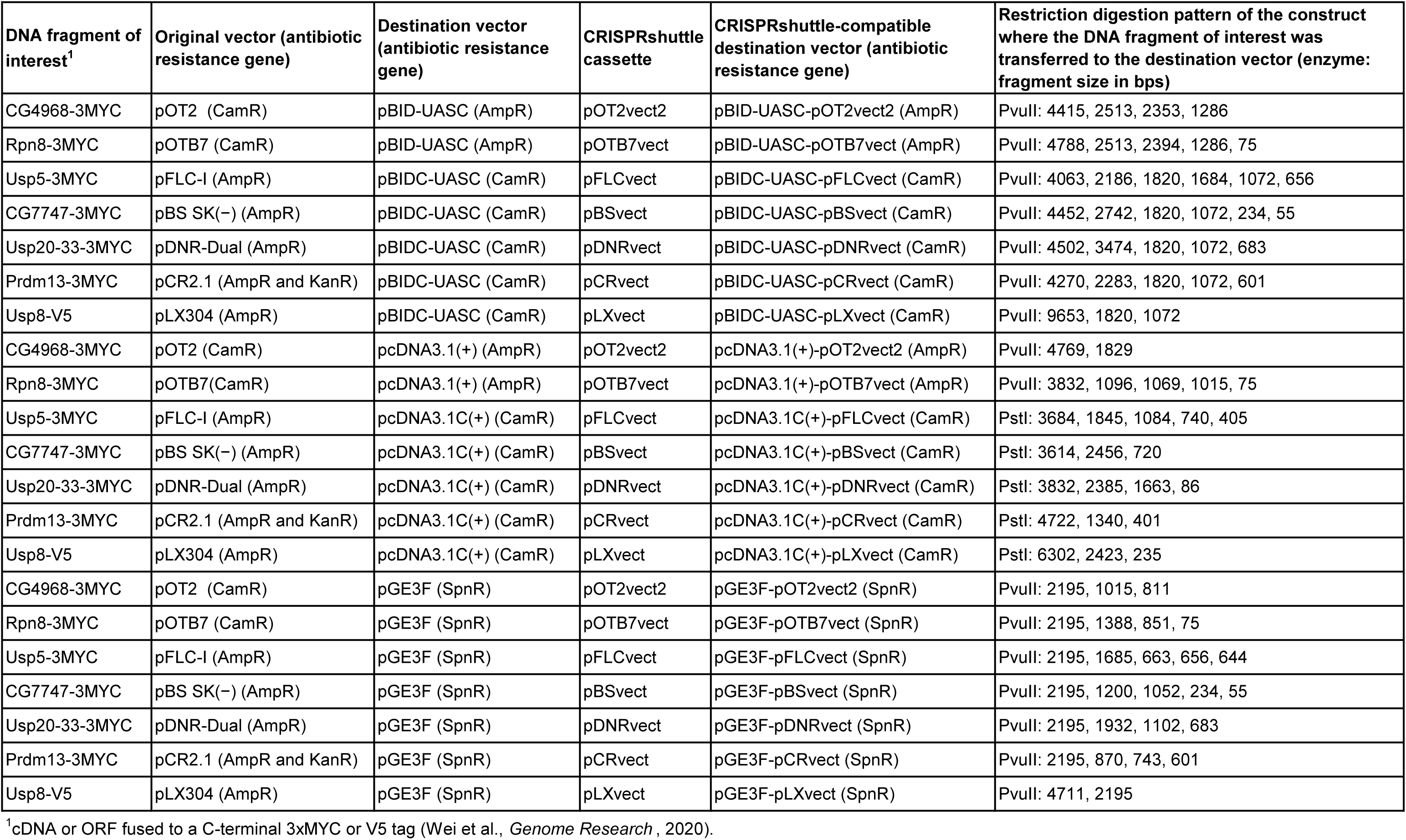
21 constructs resulting from transfer of the DNA fragments of interest from seven original vectors to three destination vectors using CRISPRshuttle.

**Supplementary Table S5.**
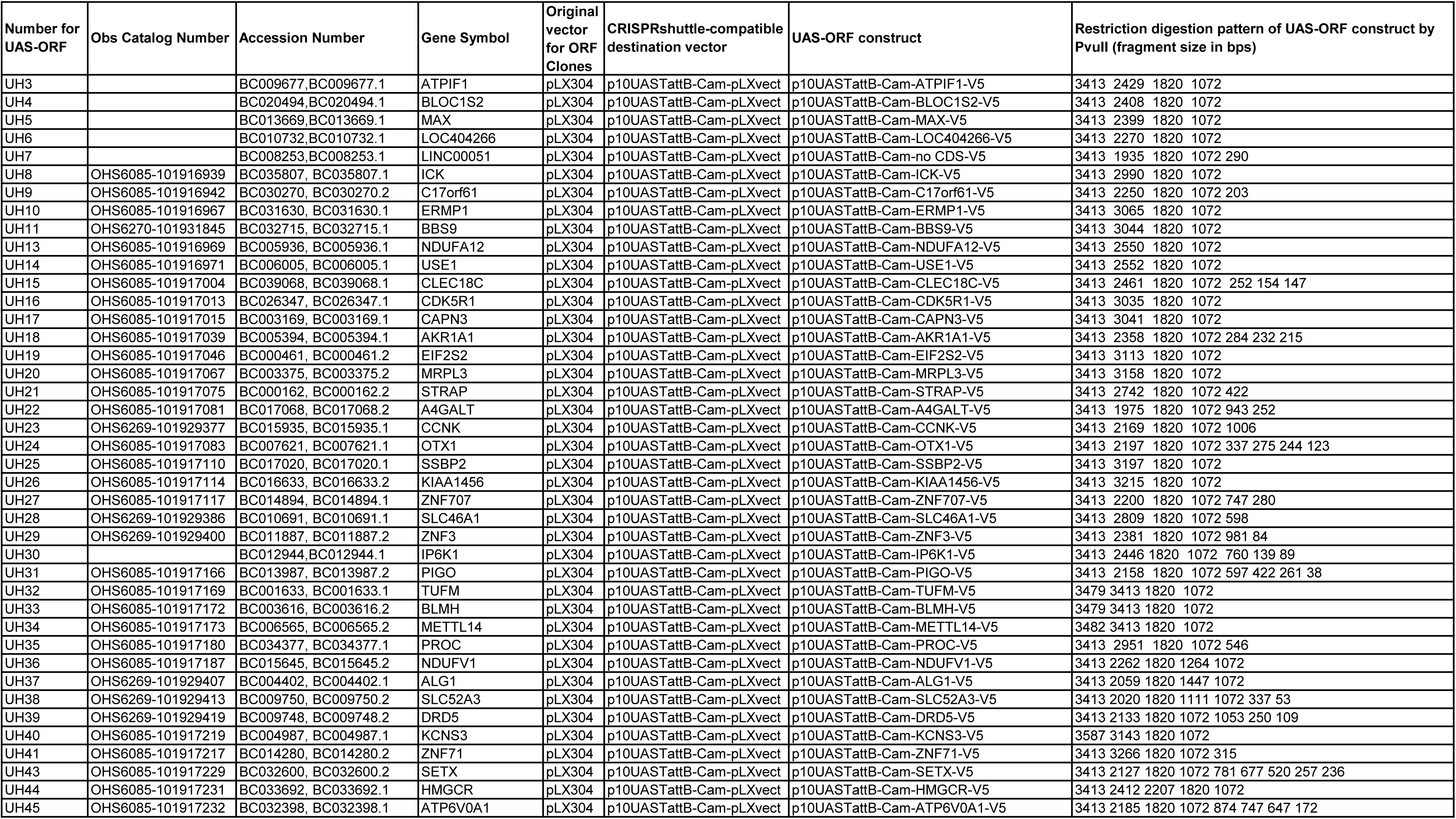
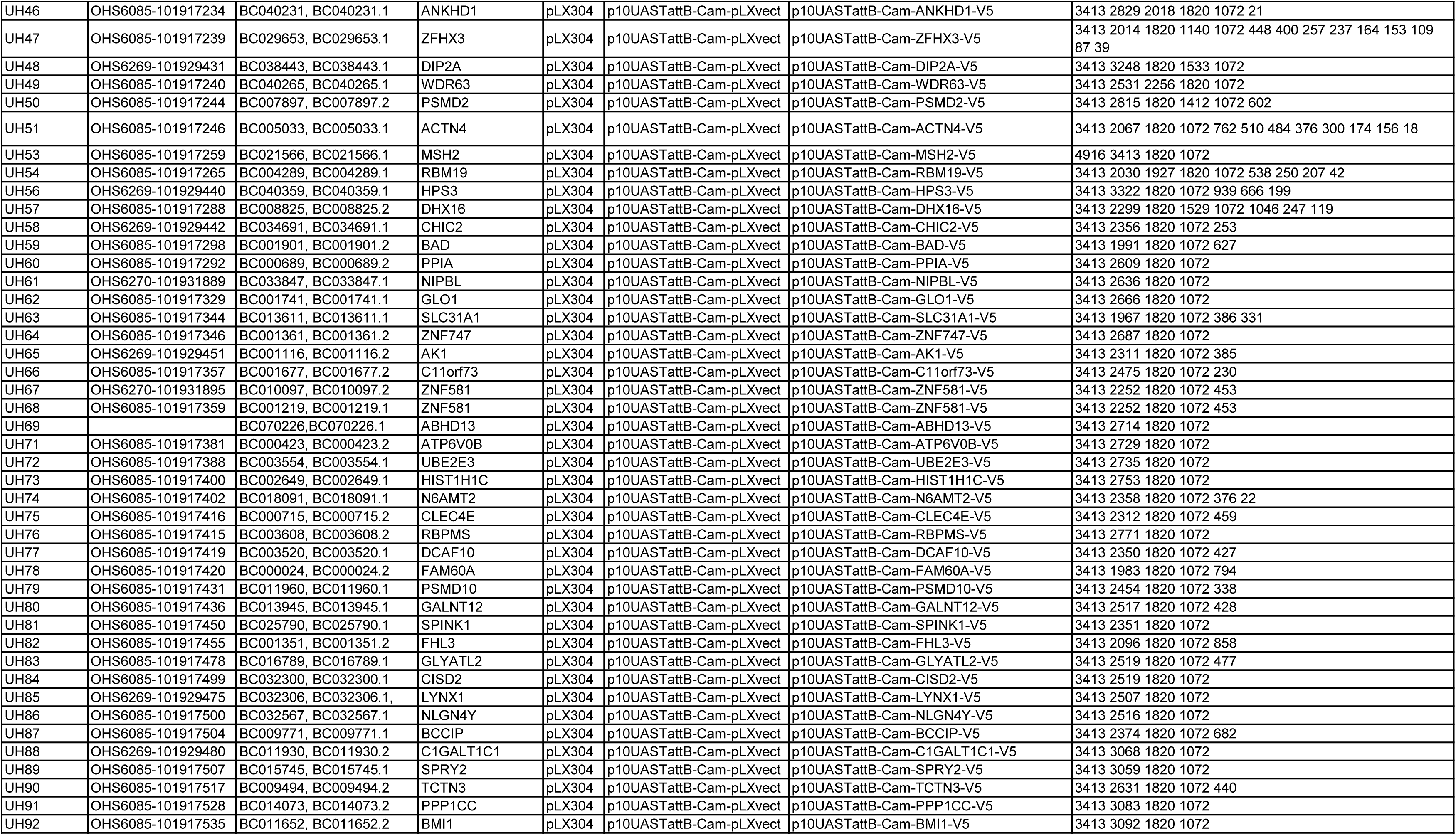
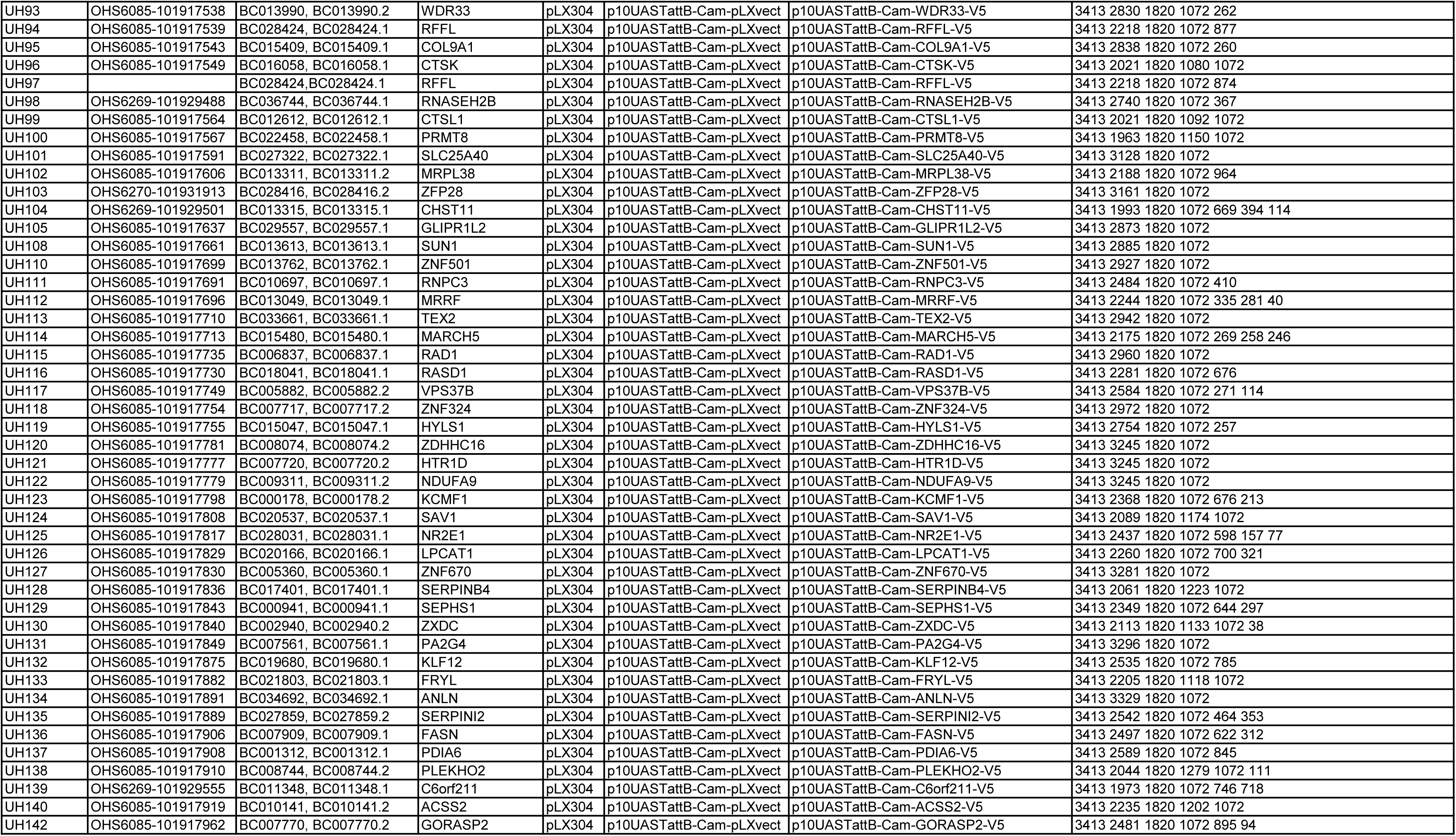
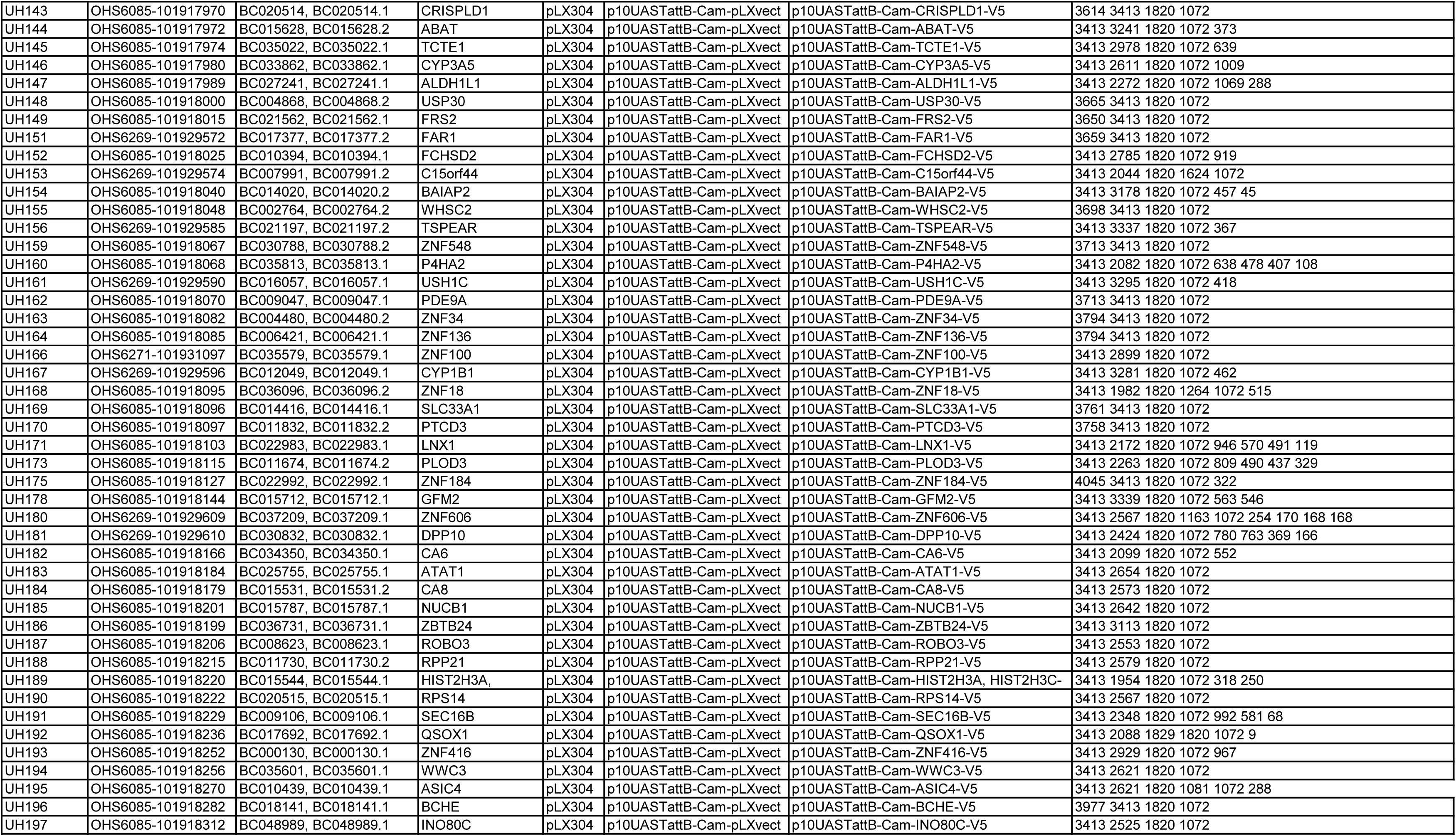
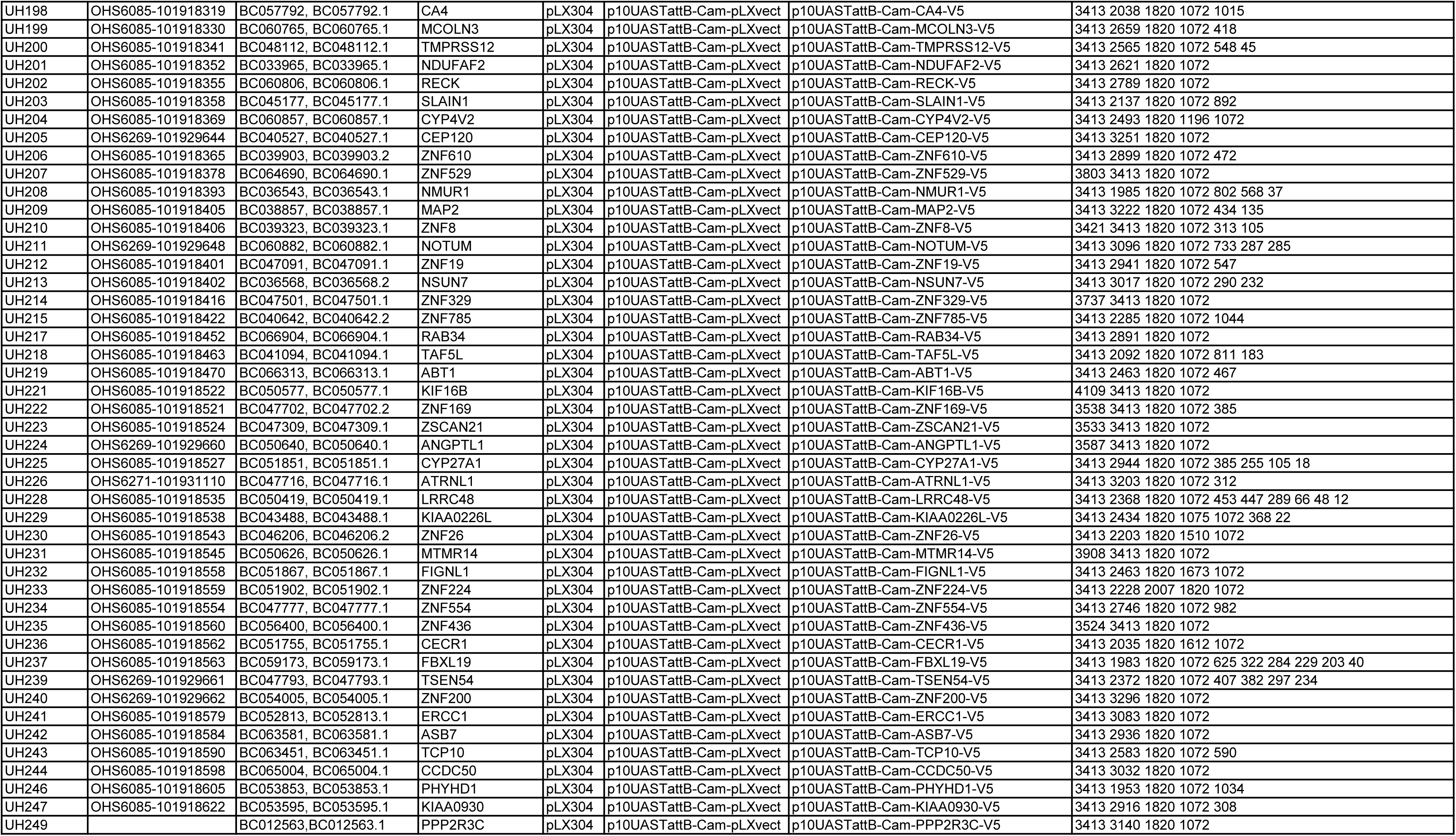
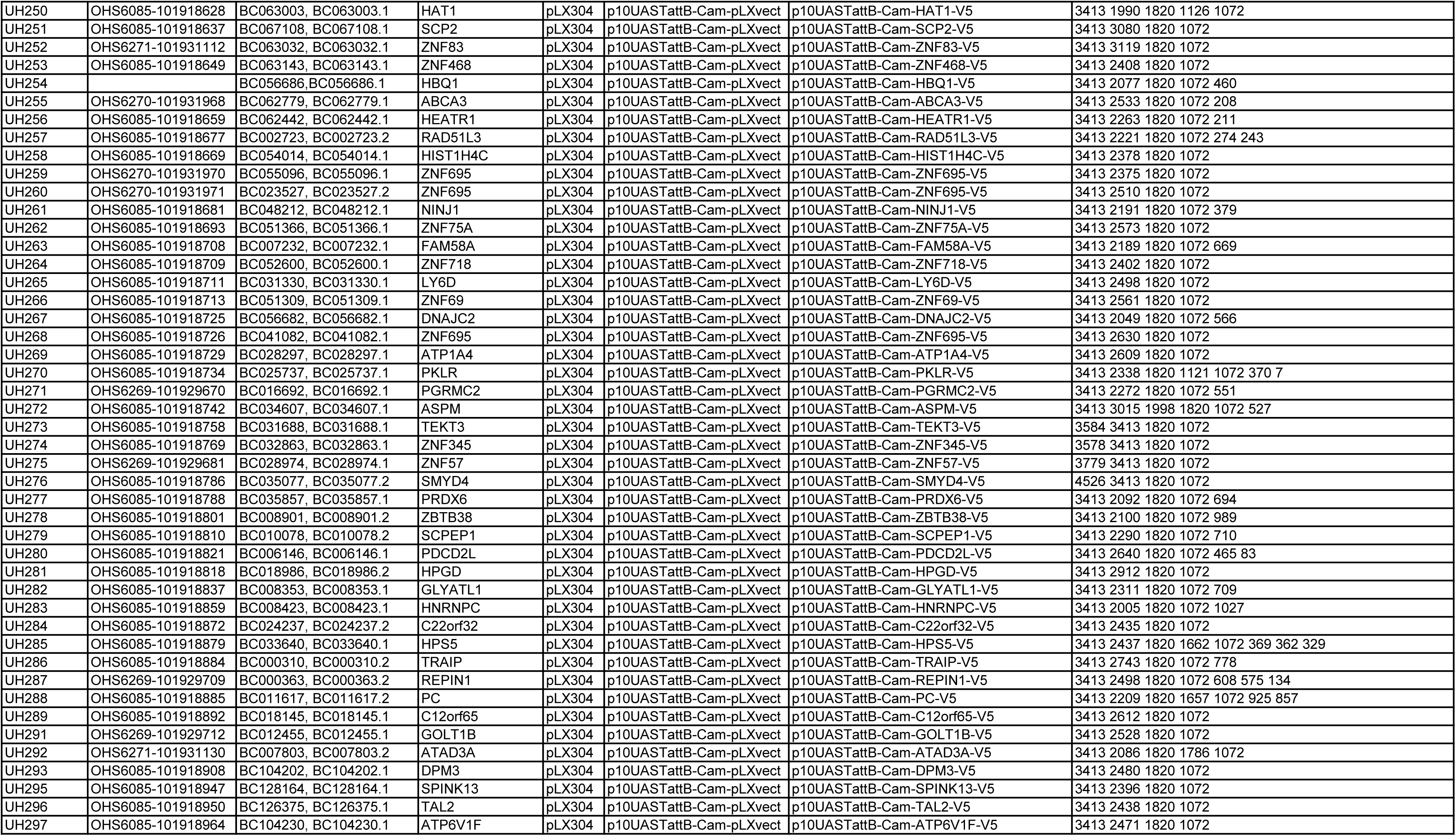
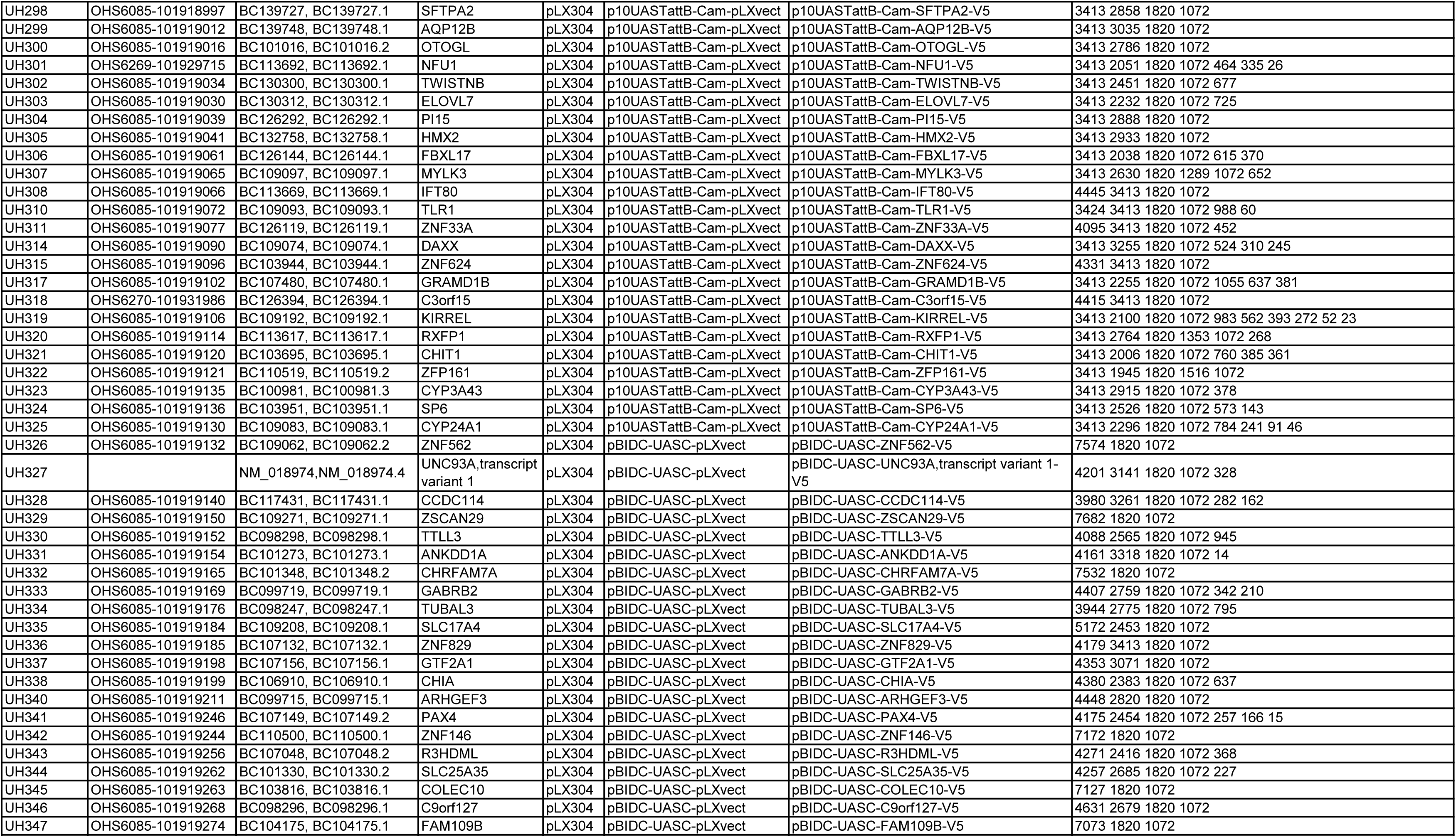
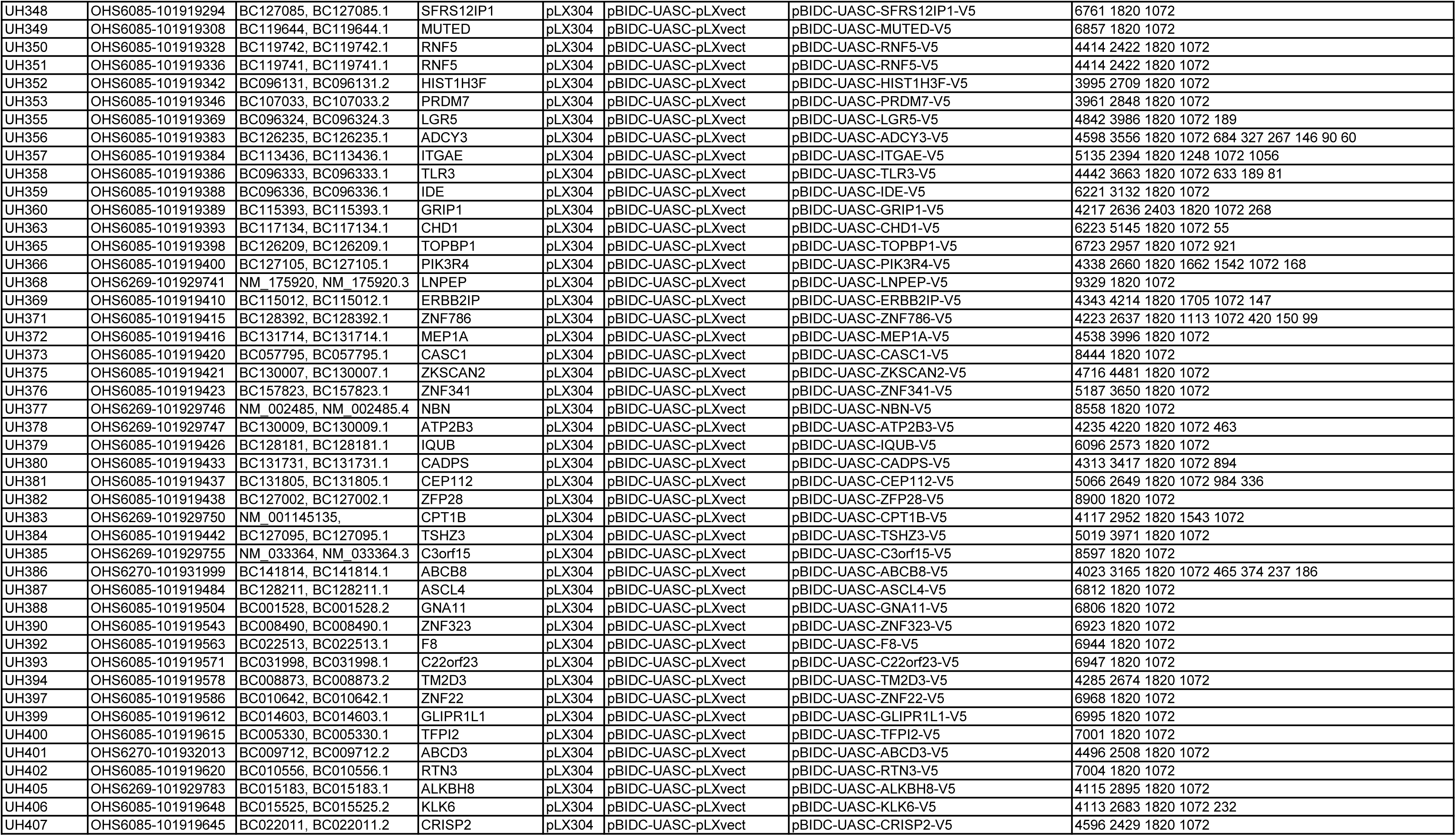
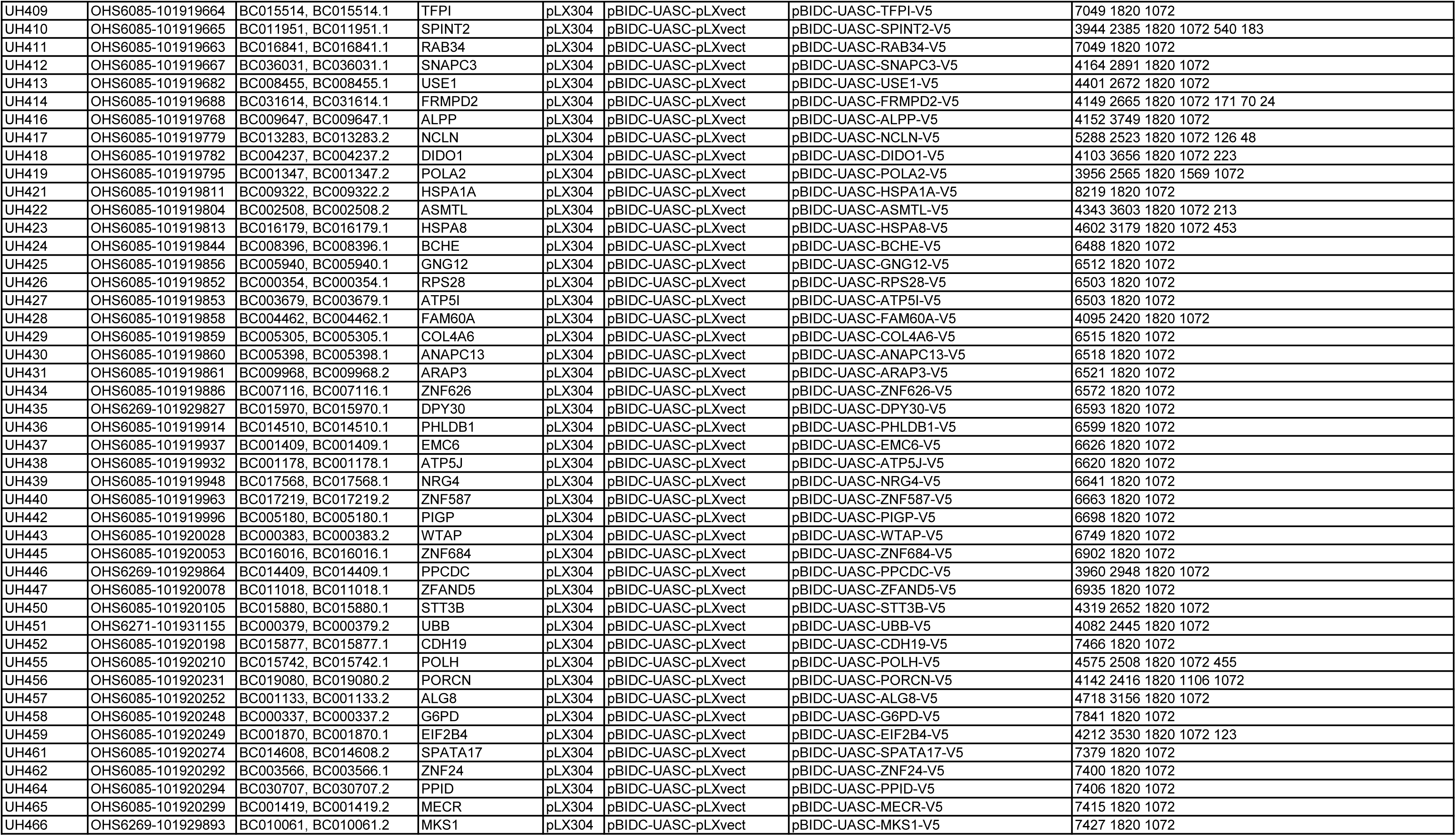
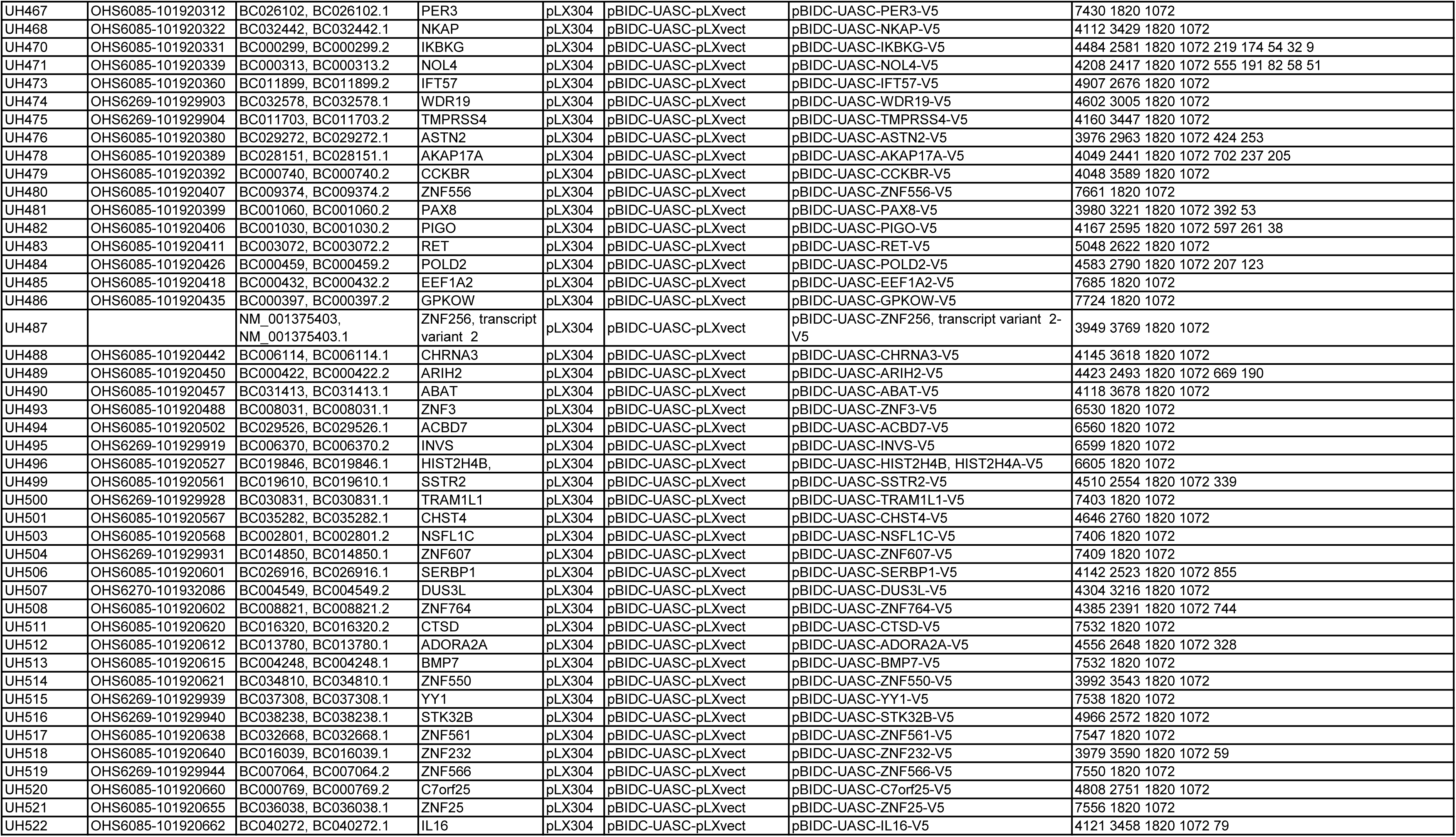
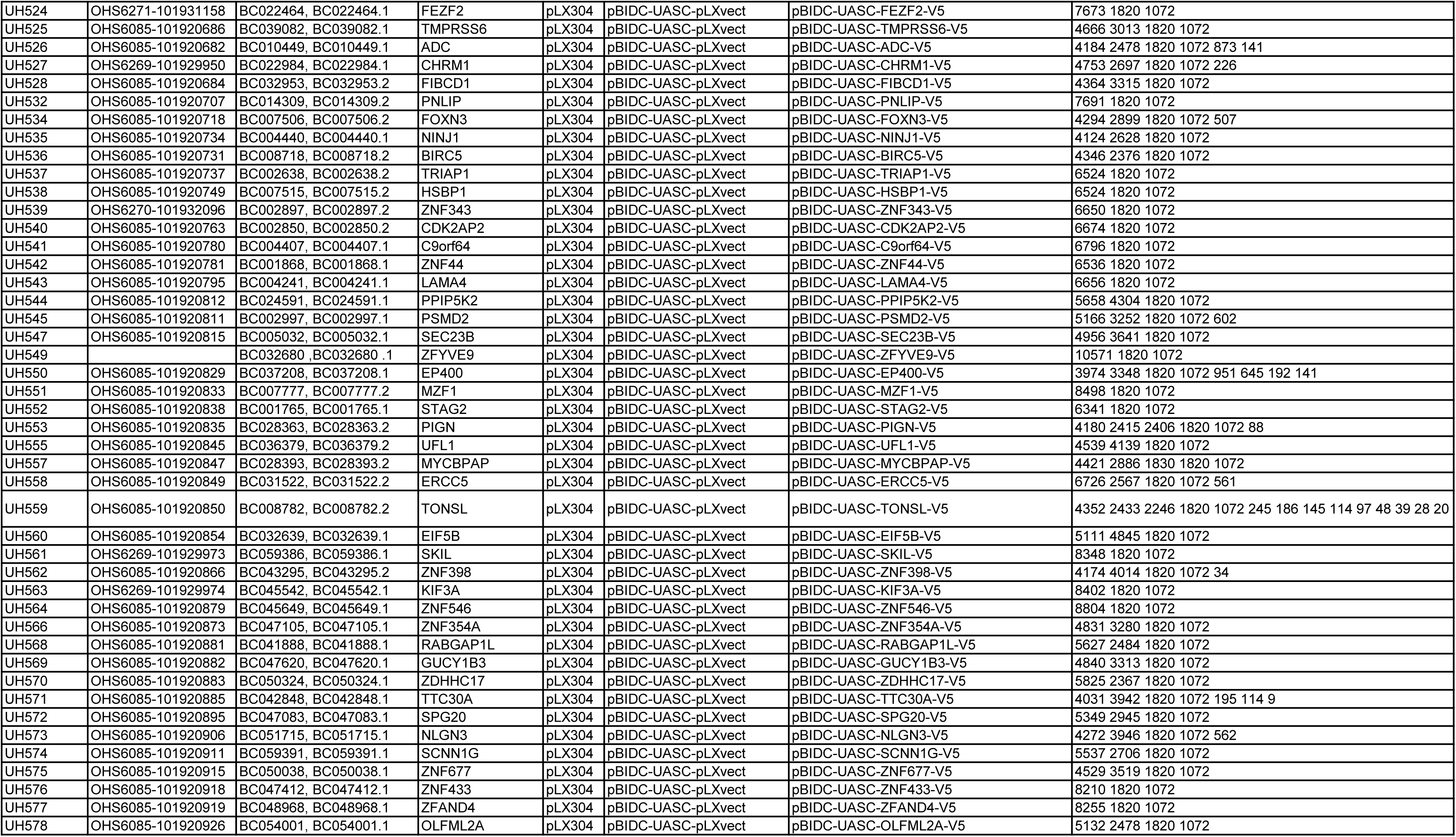
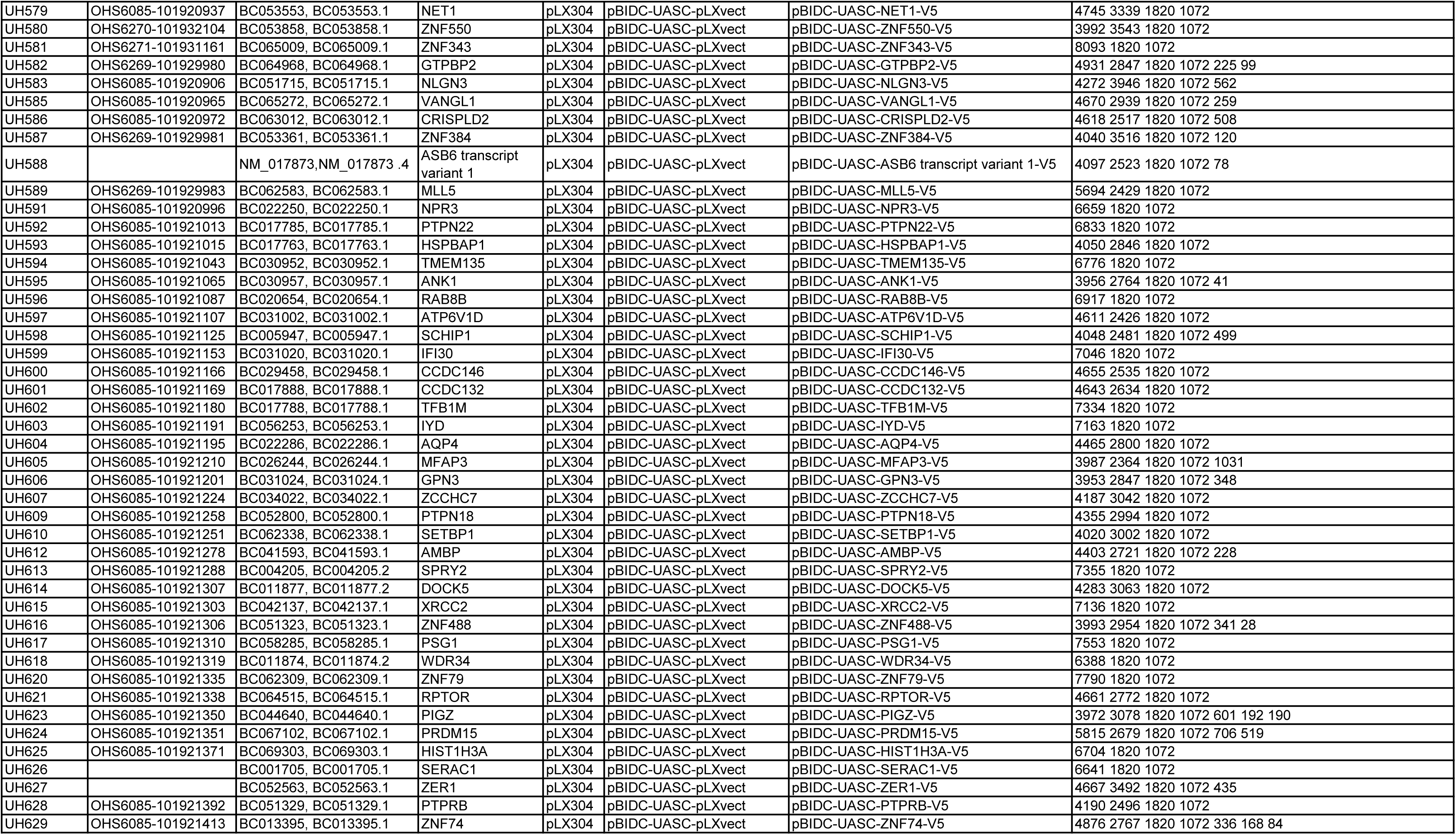
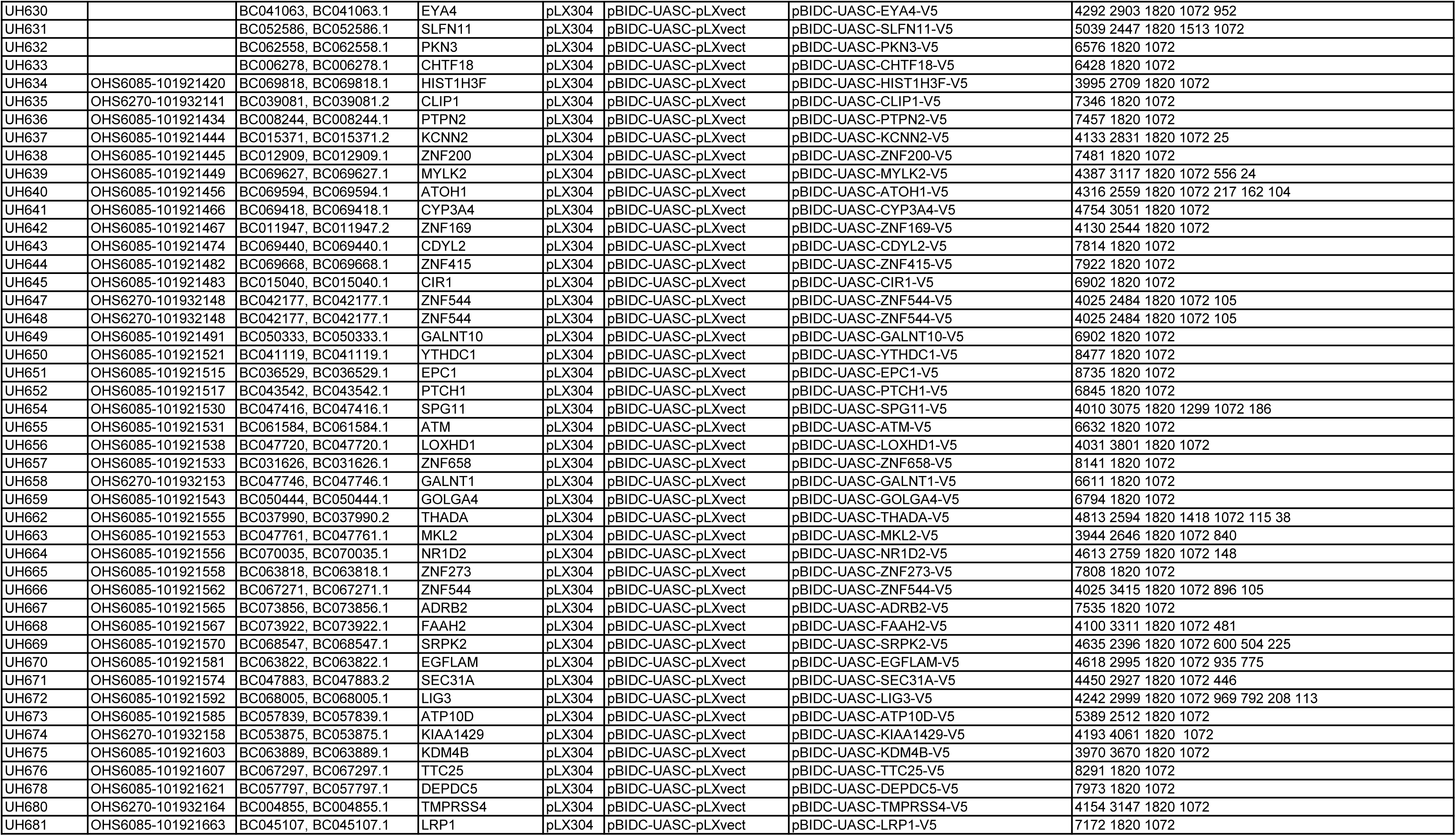
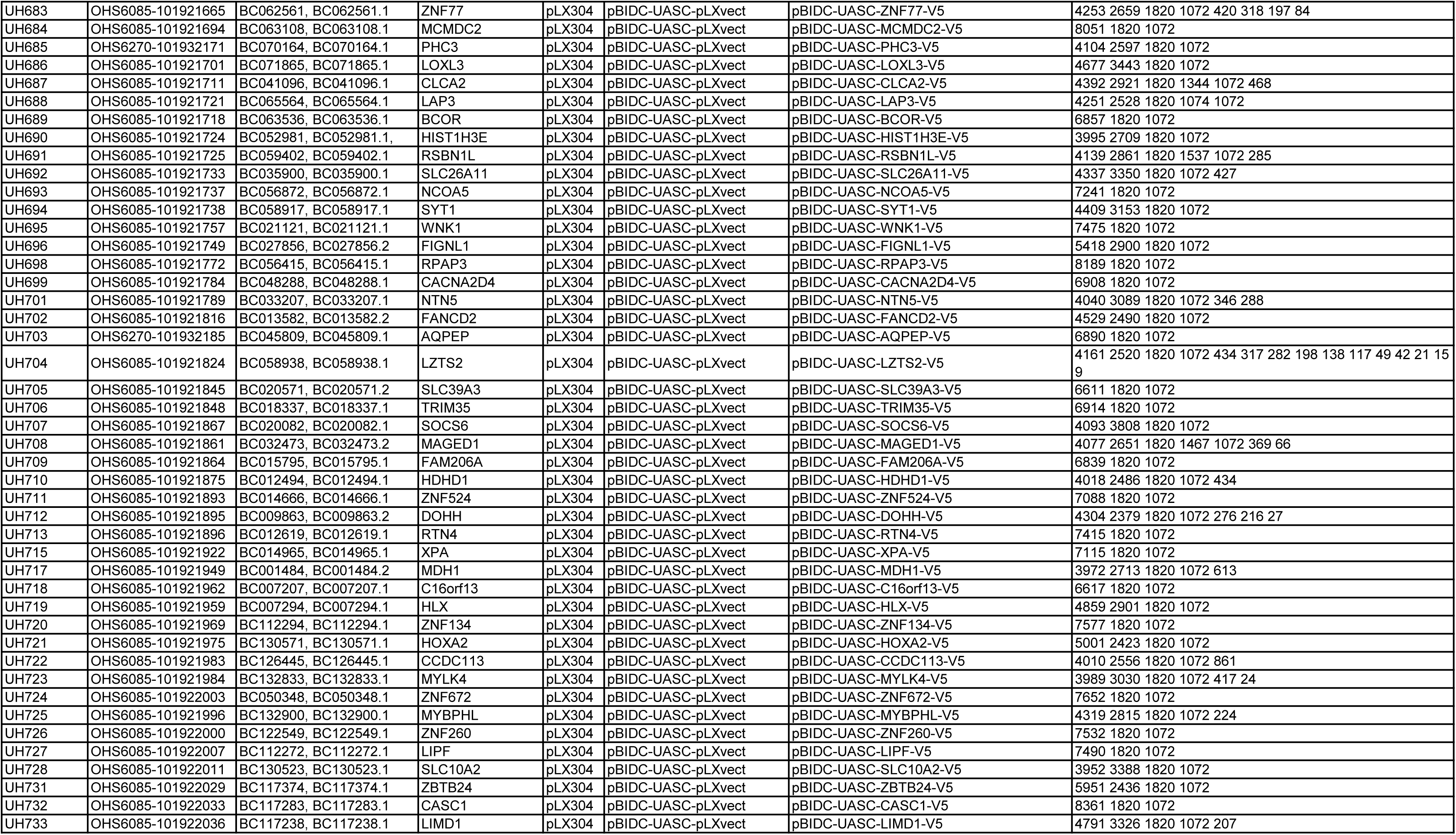
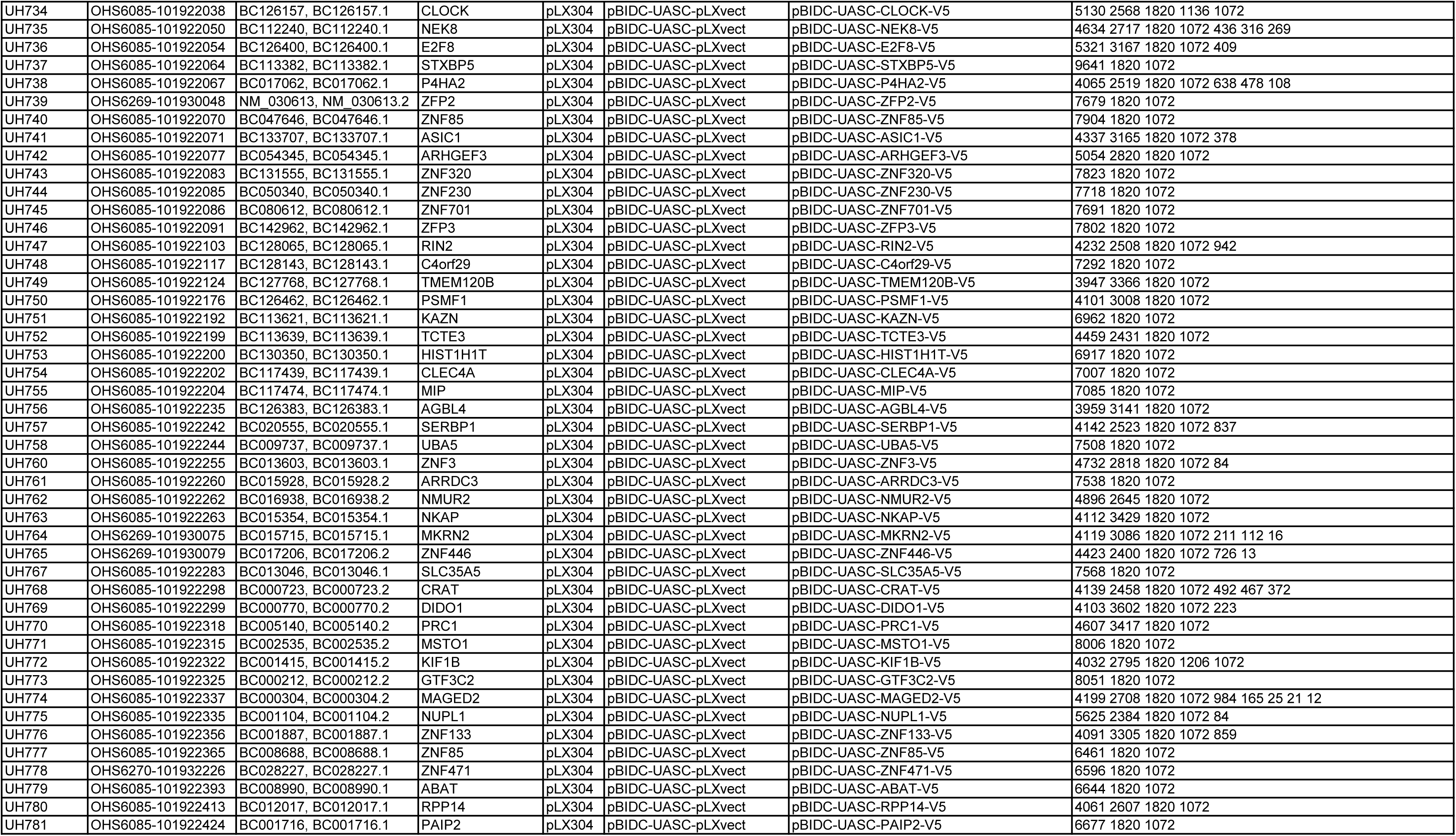
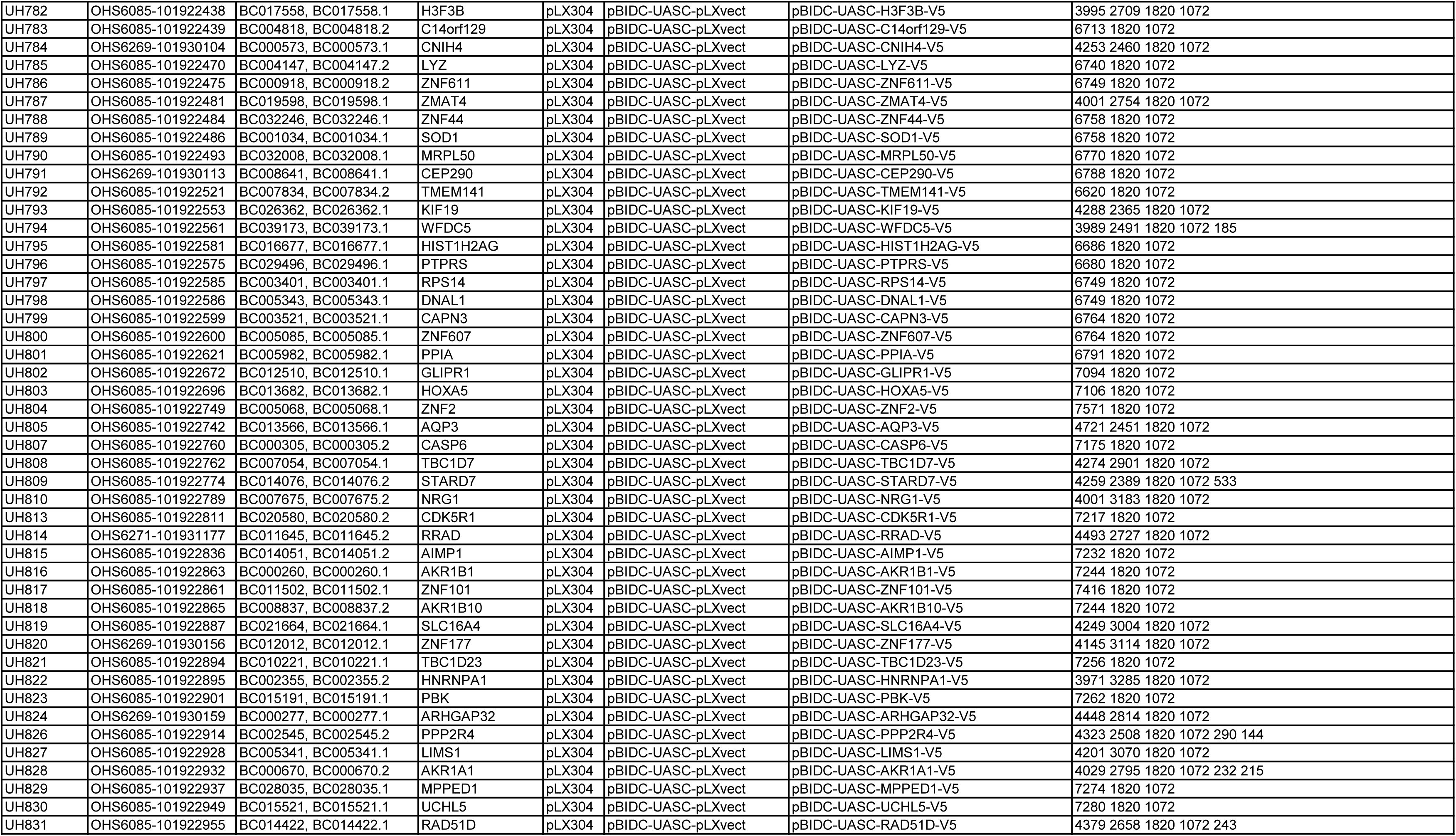
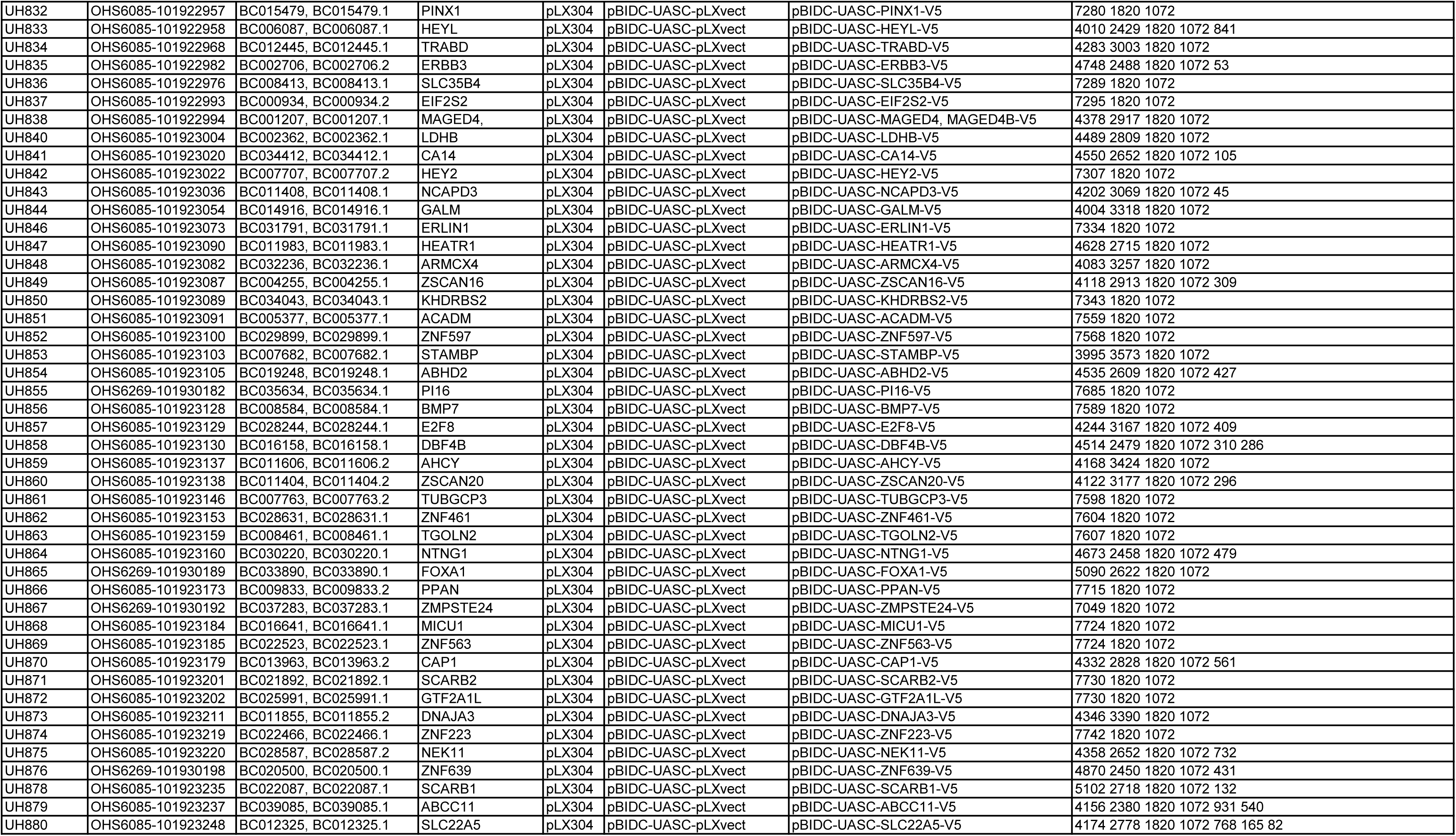
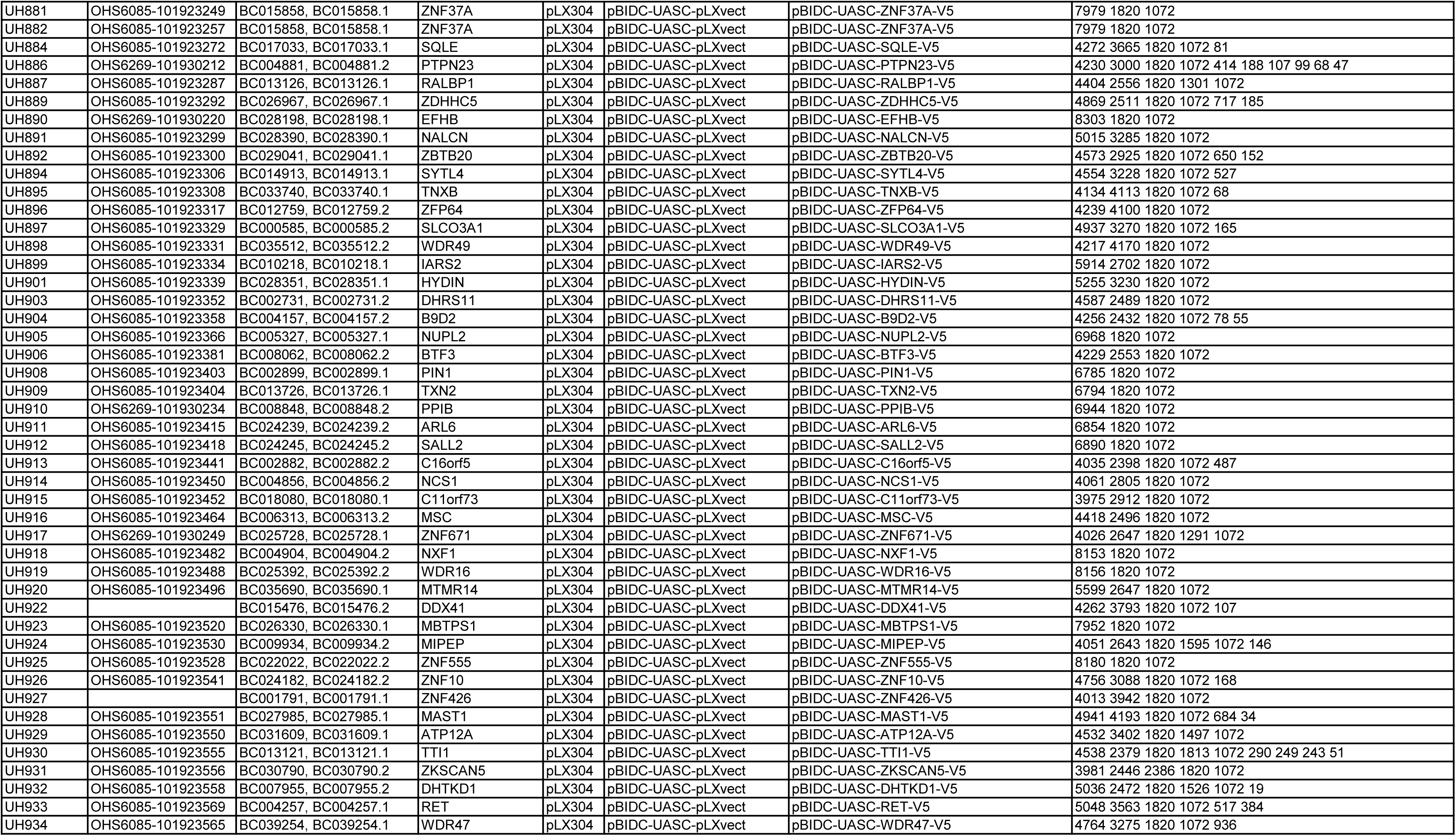
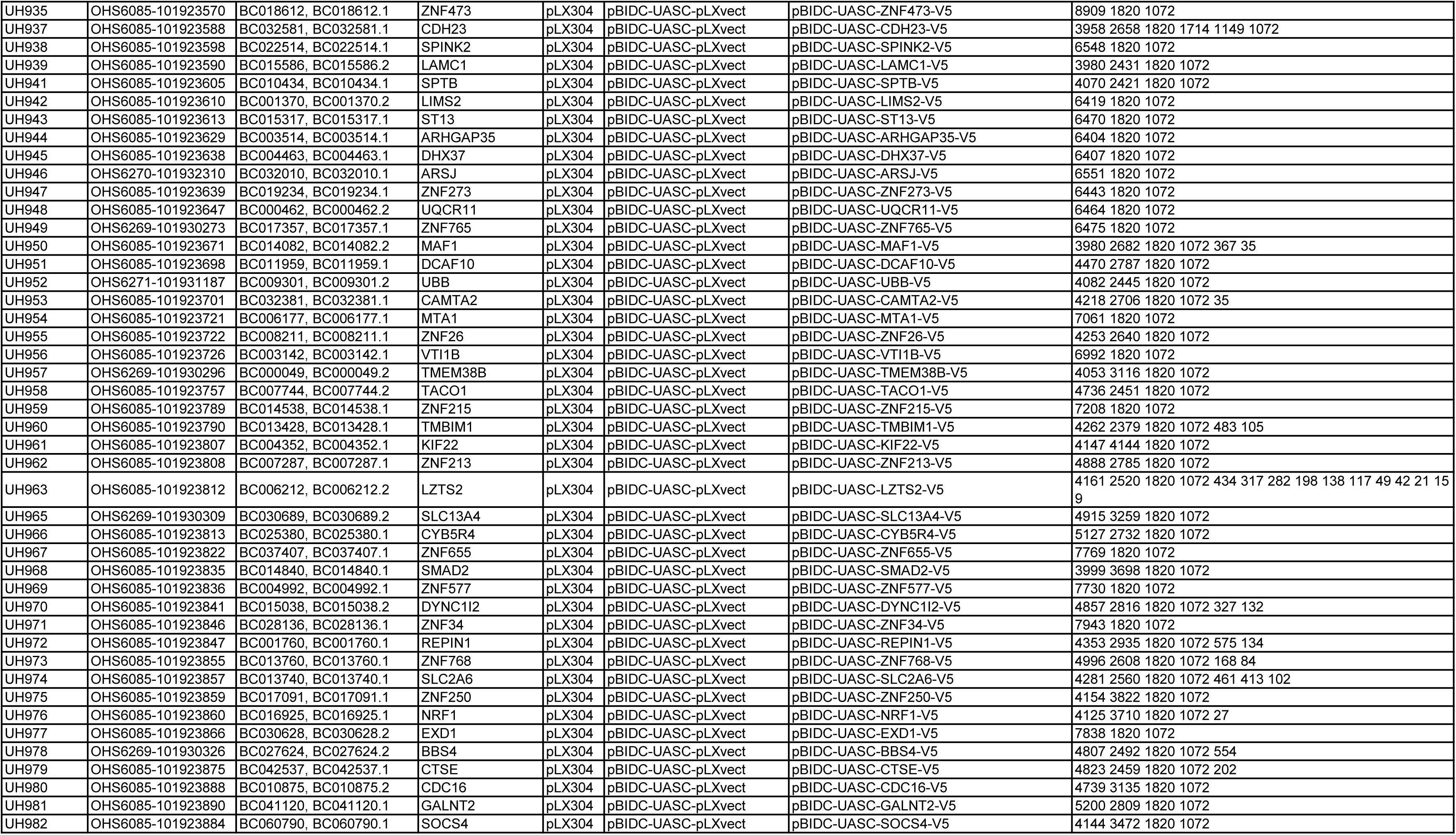
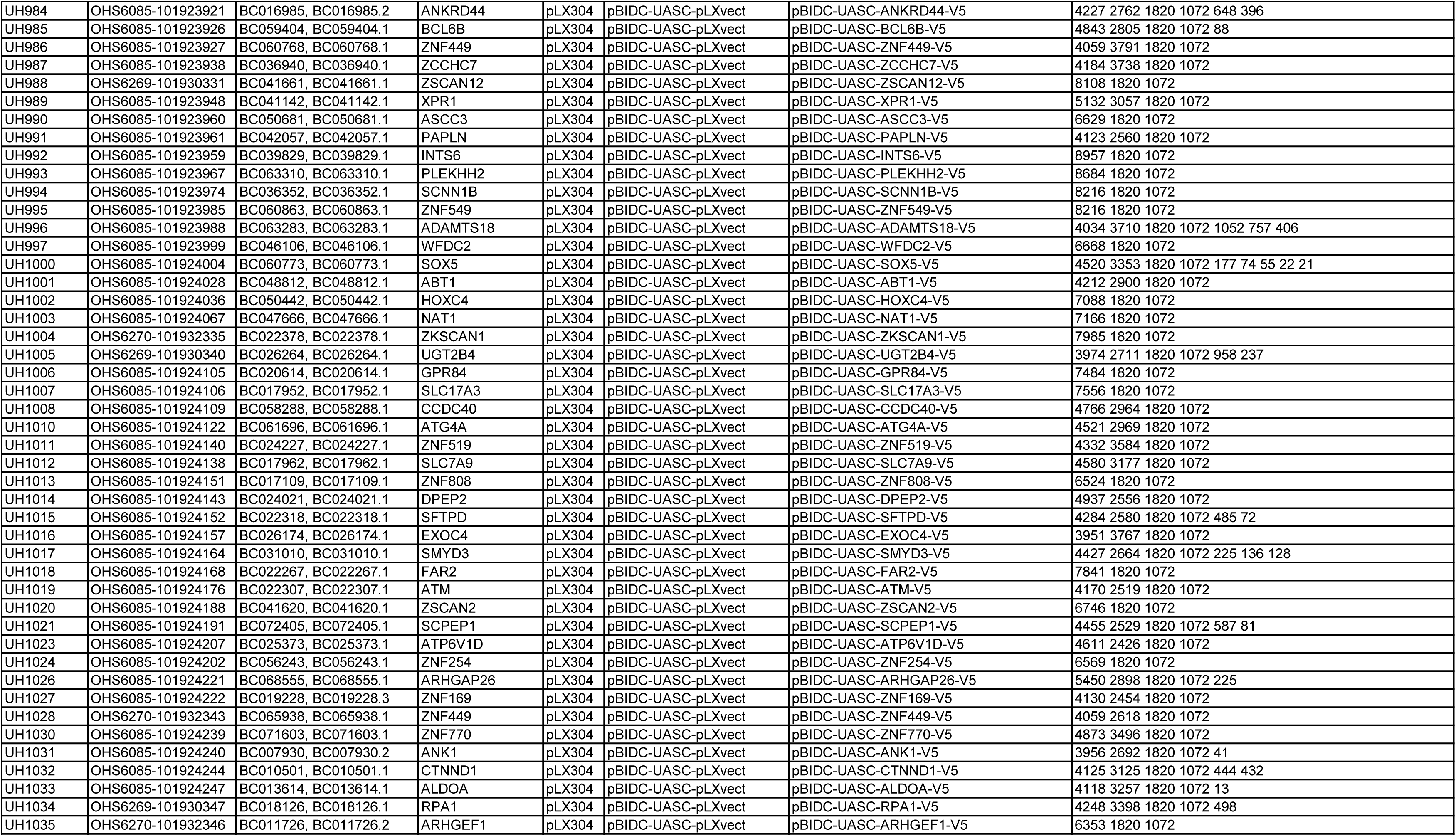
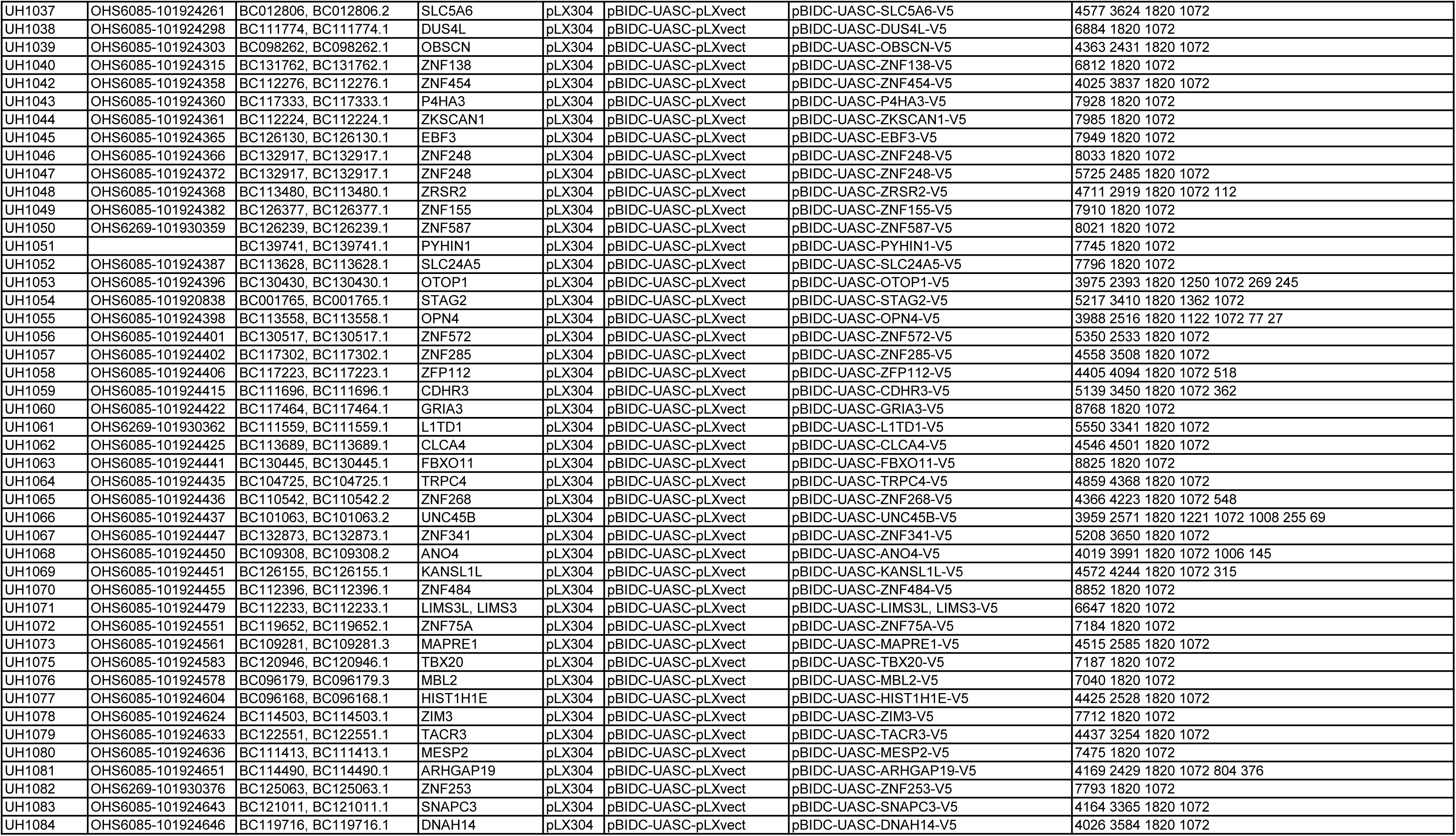
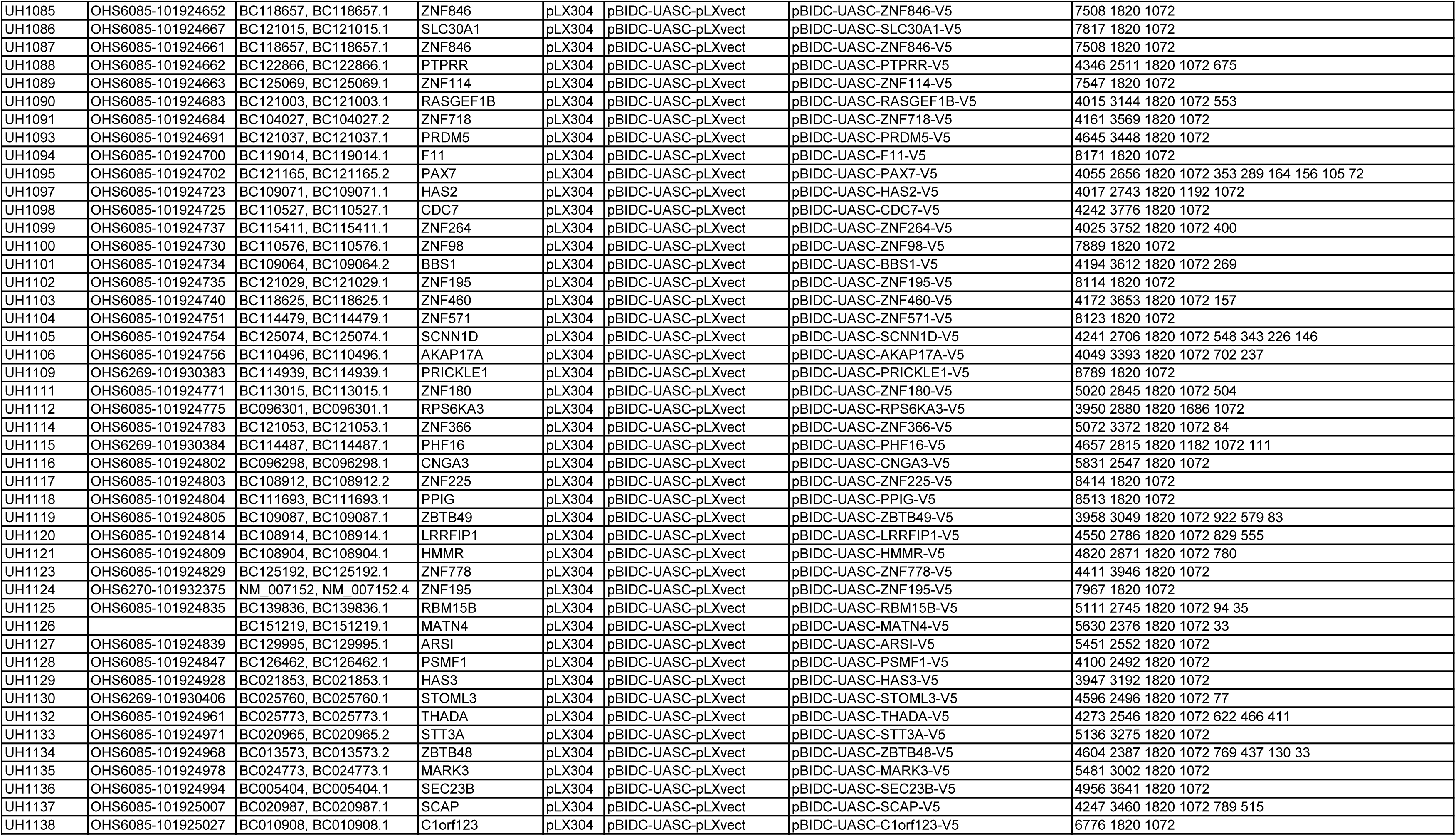
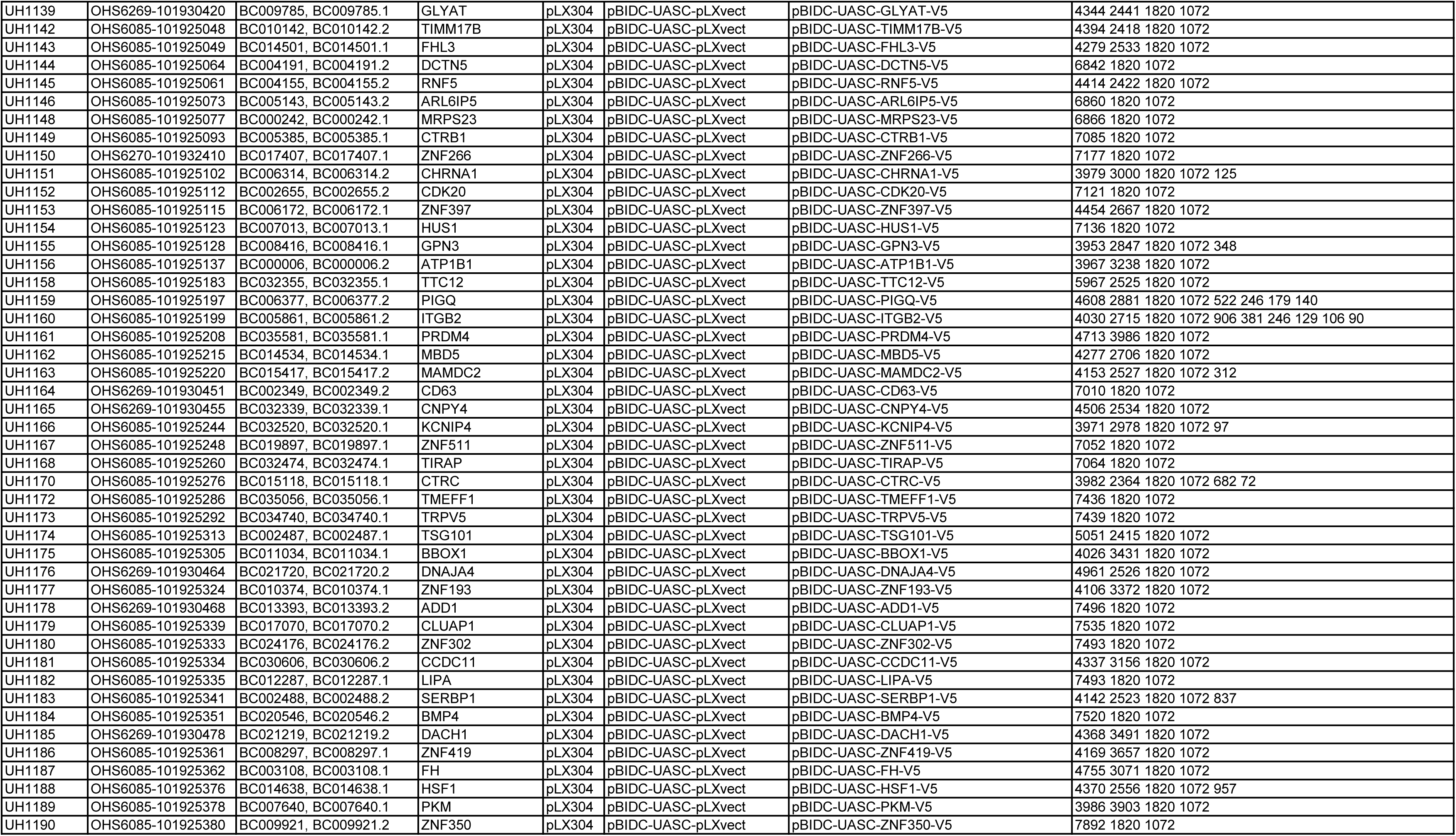
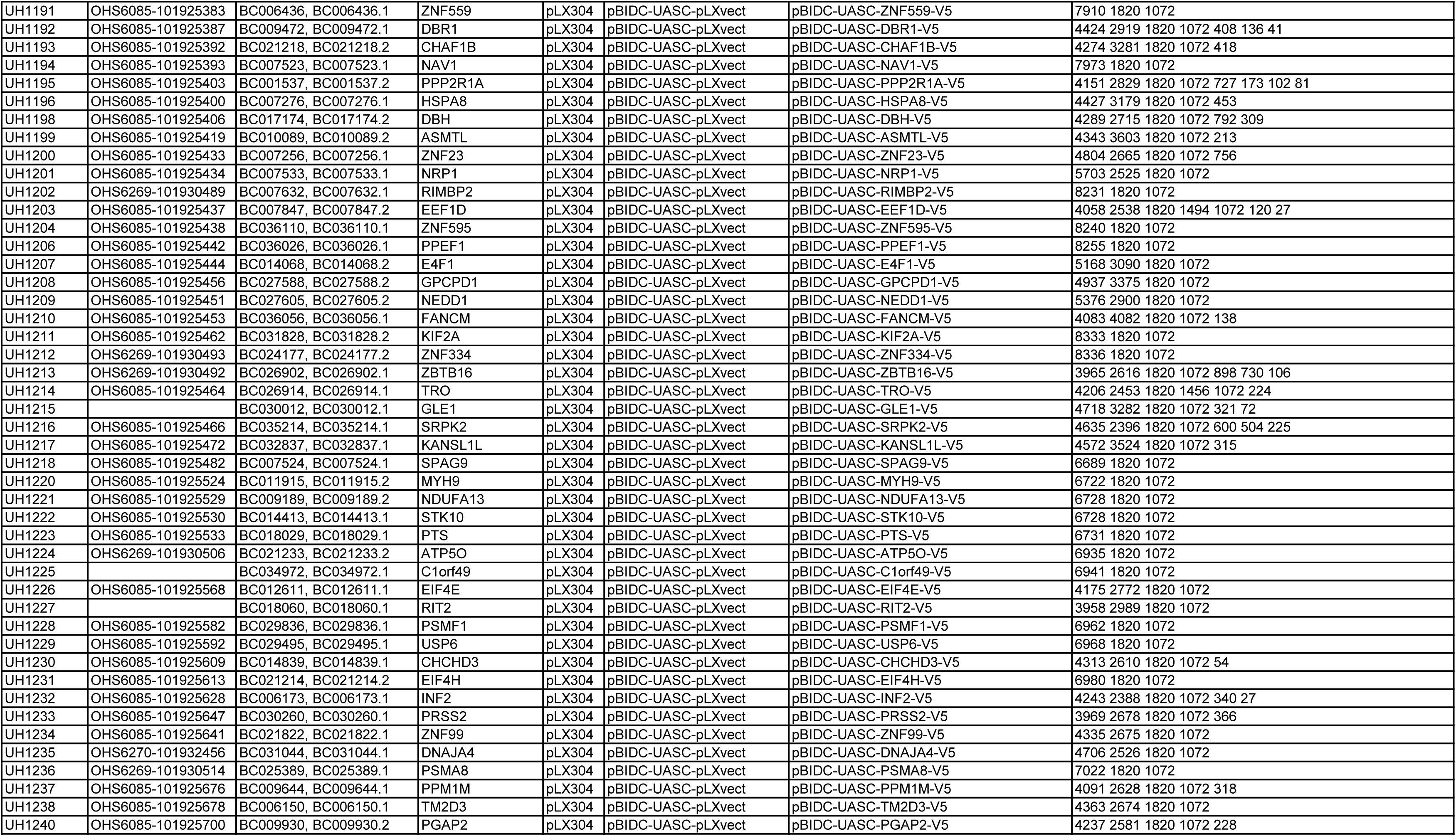
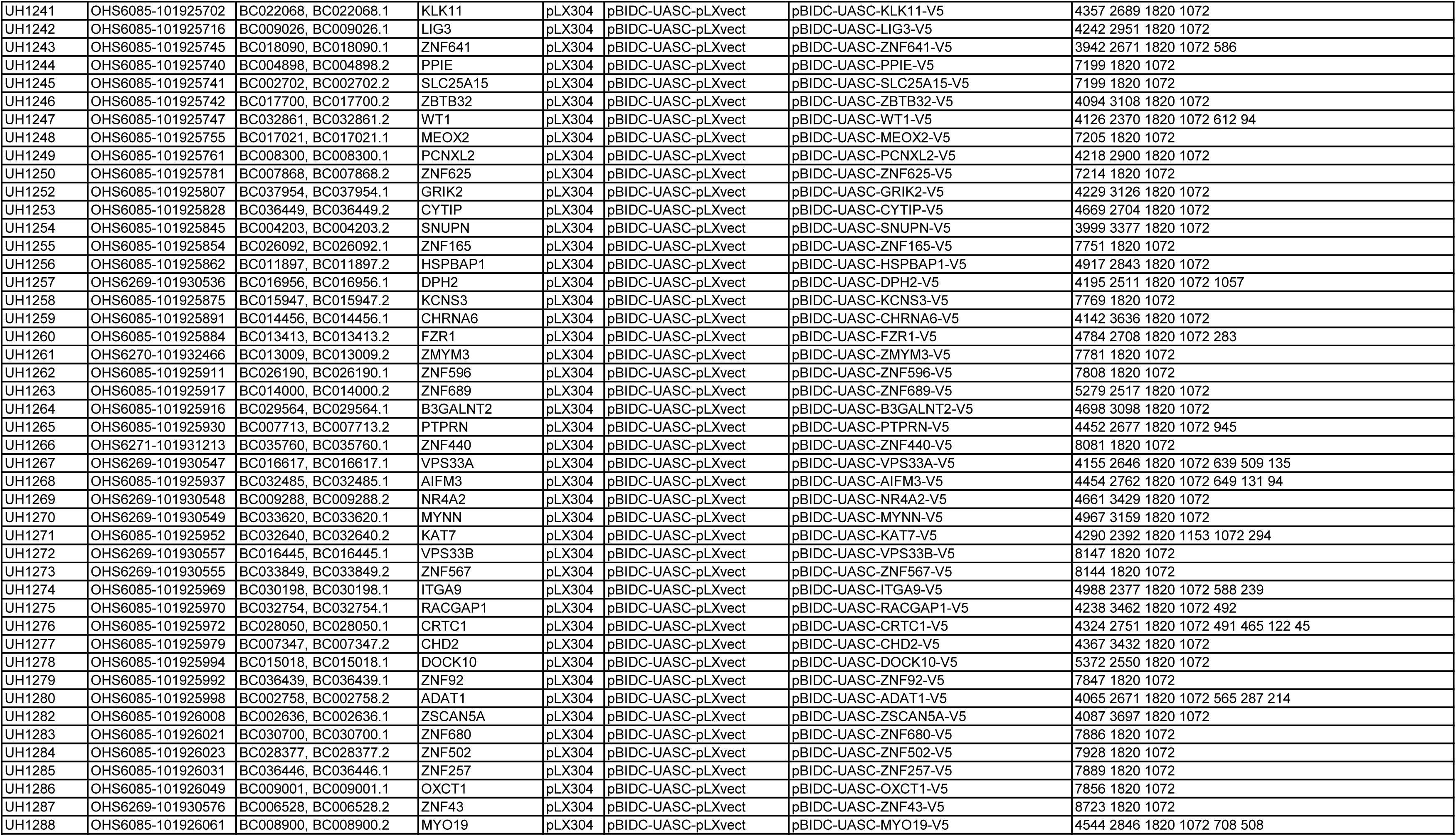
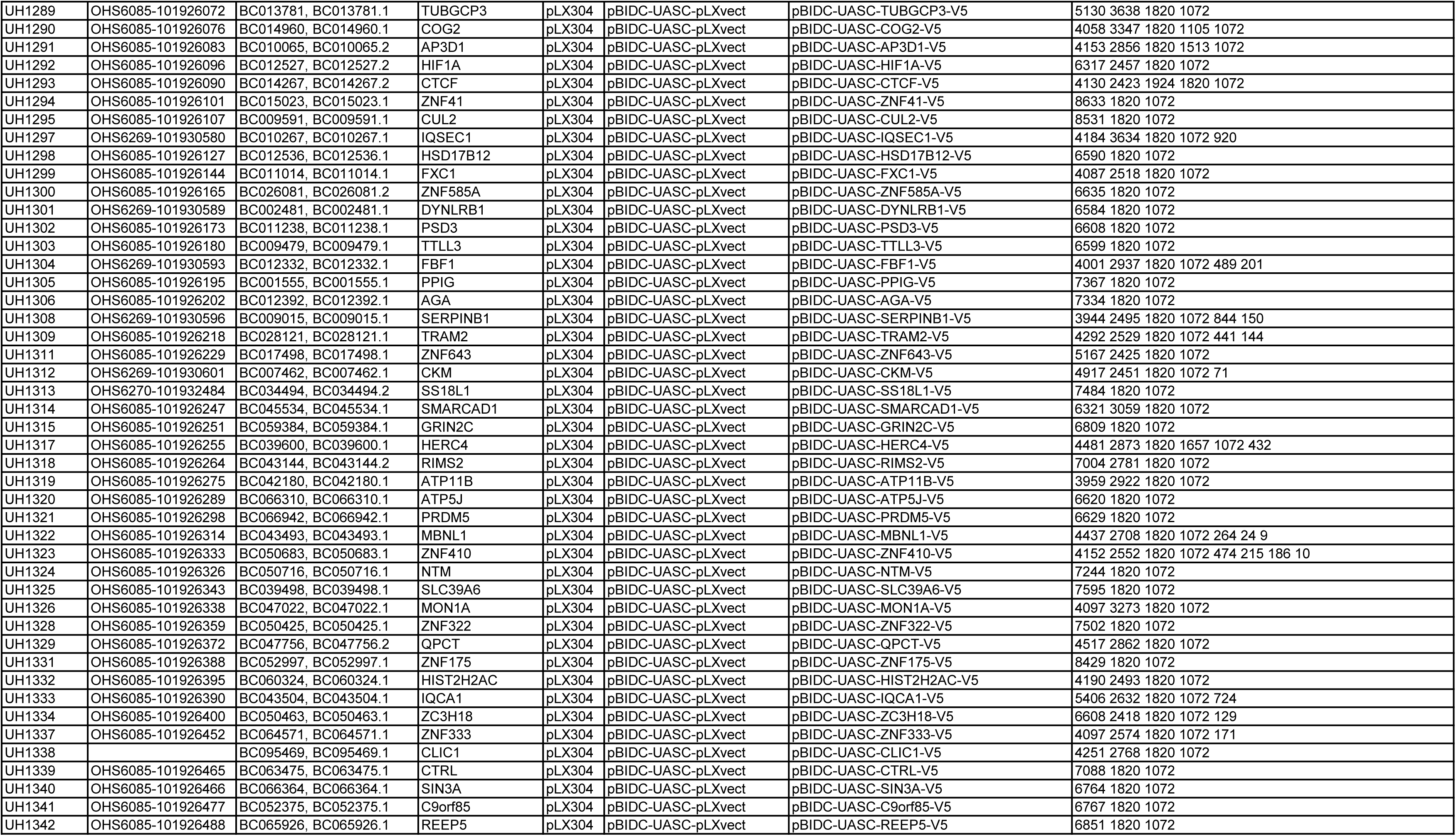
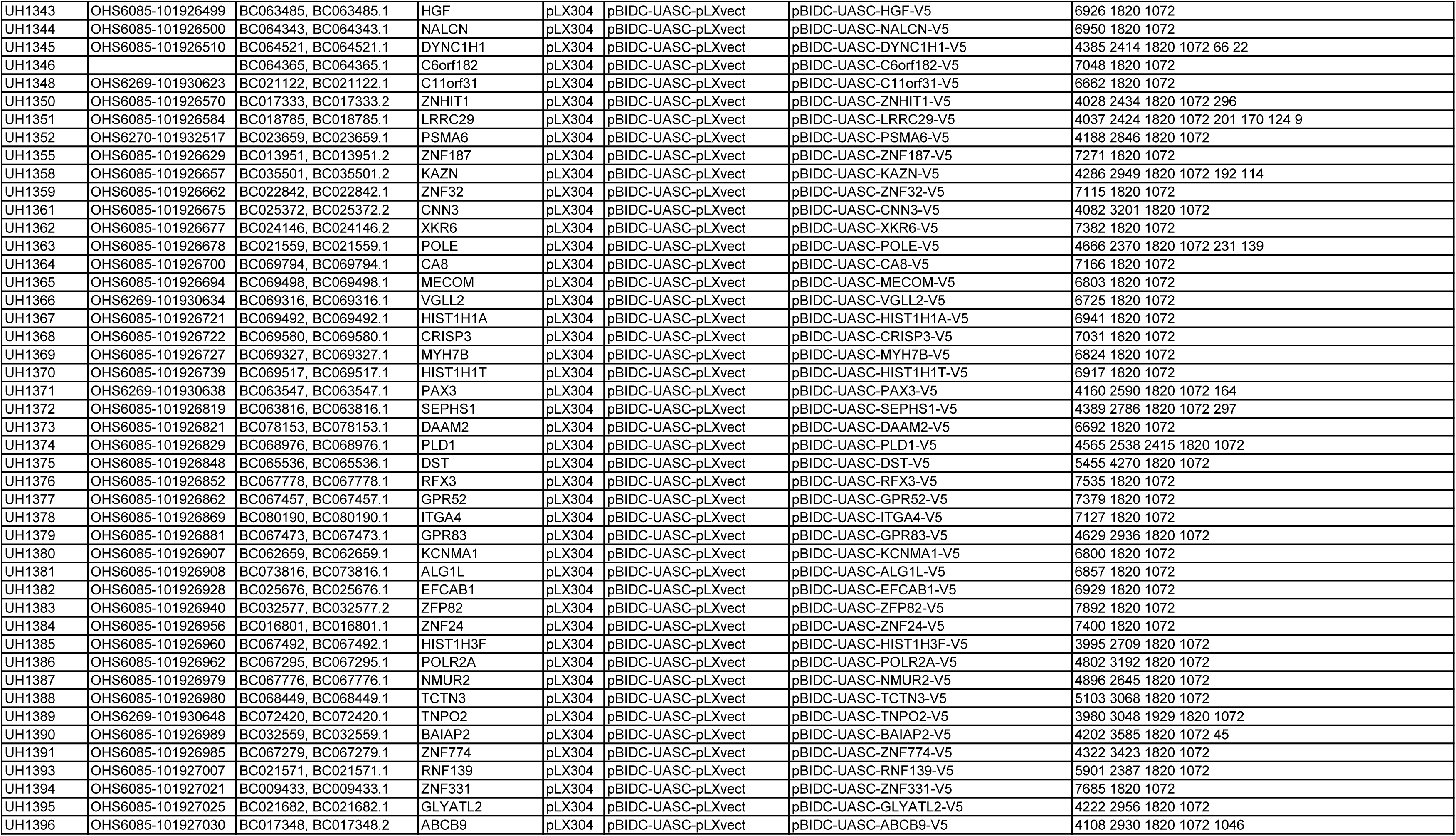
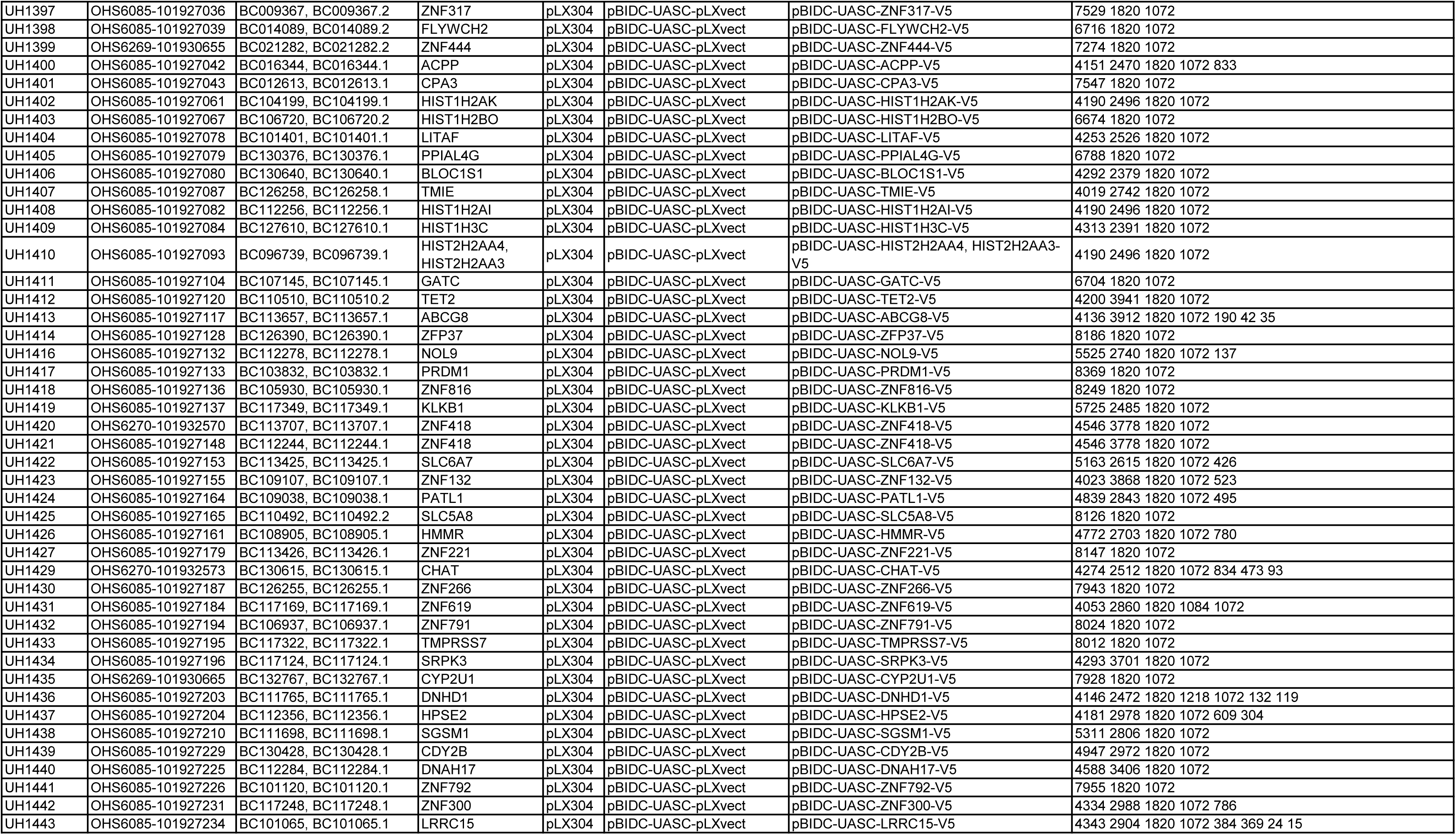
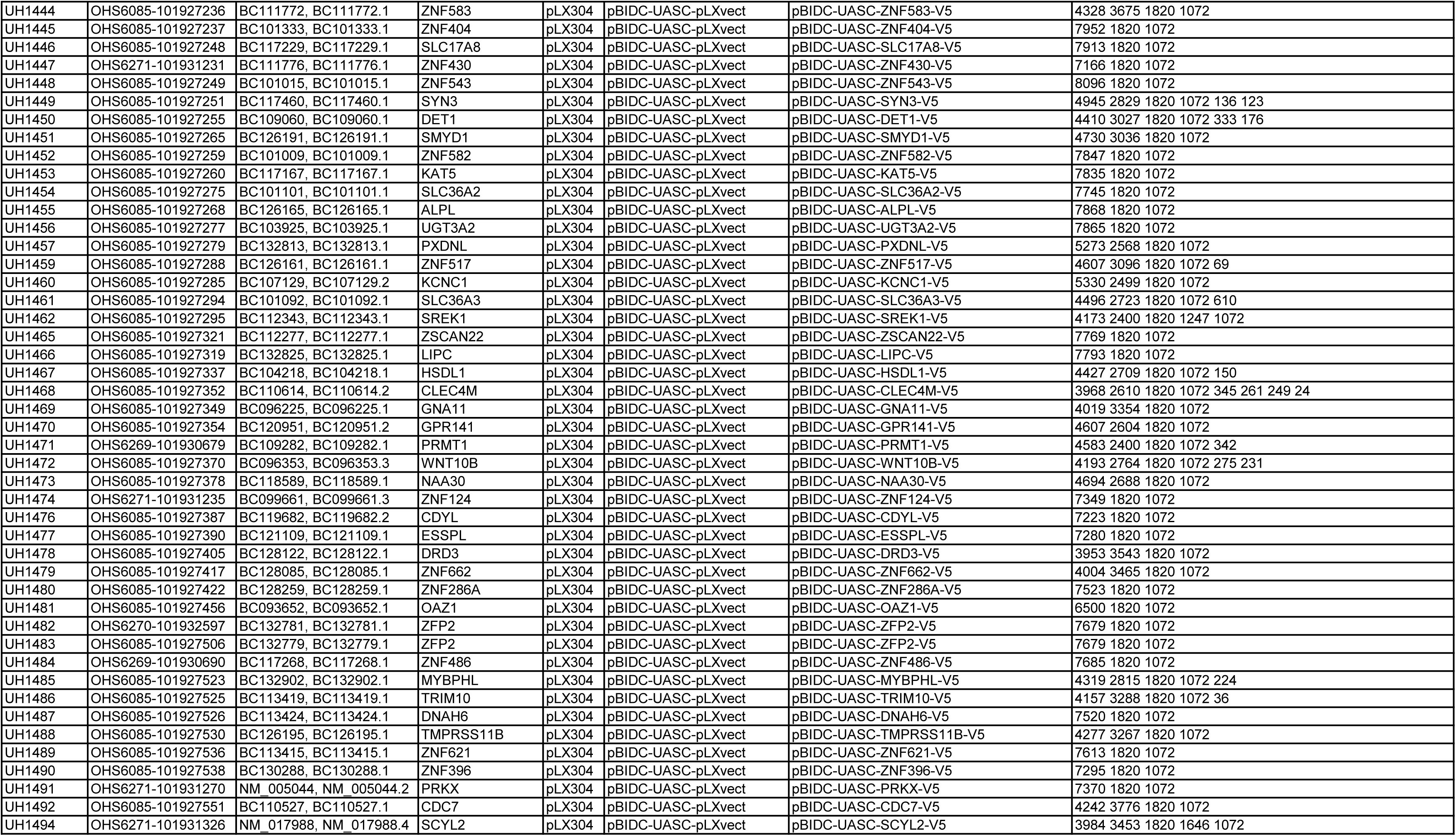
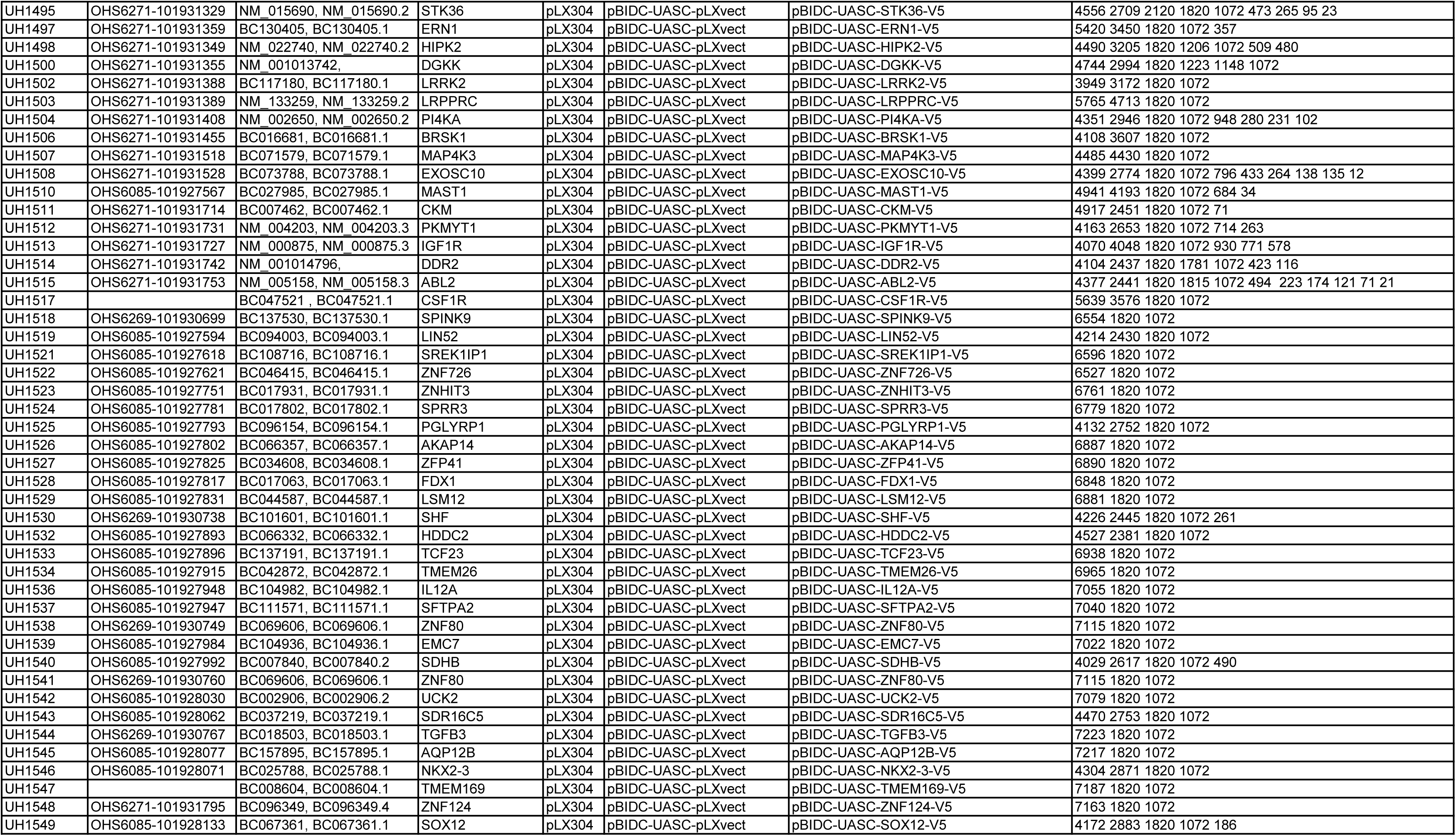
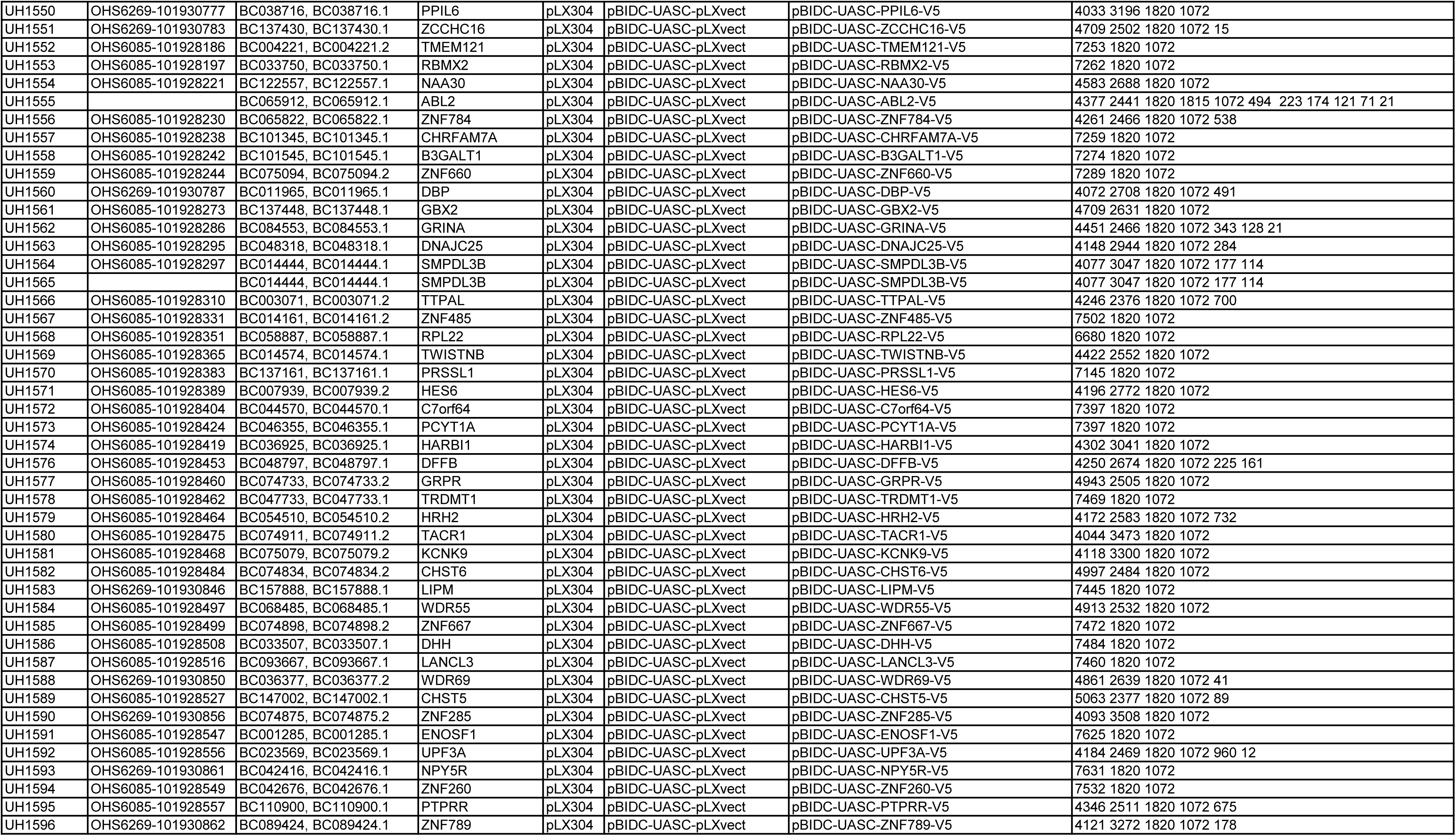
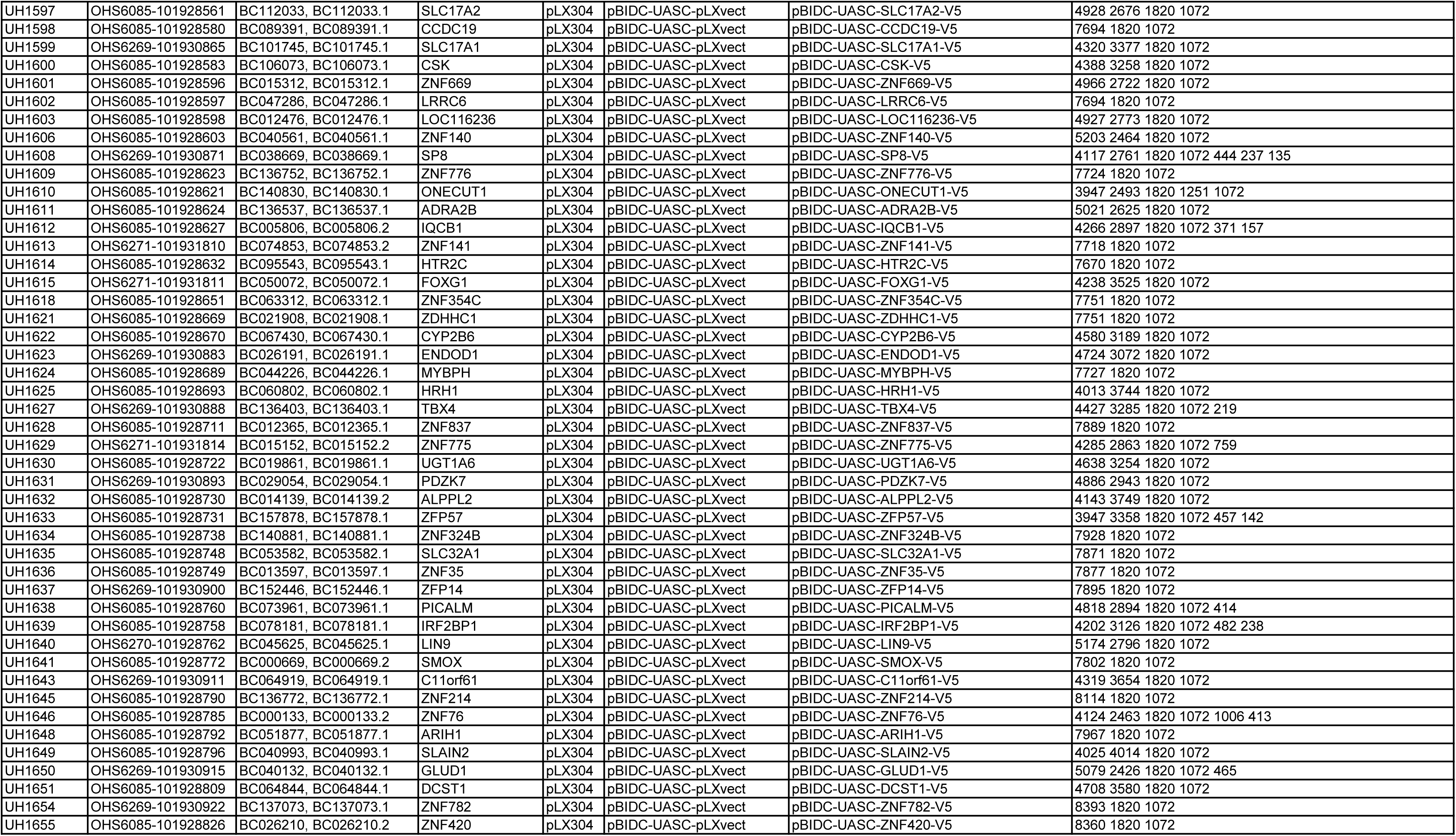
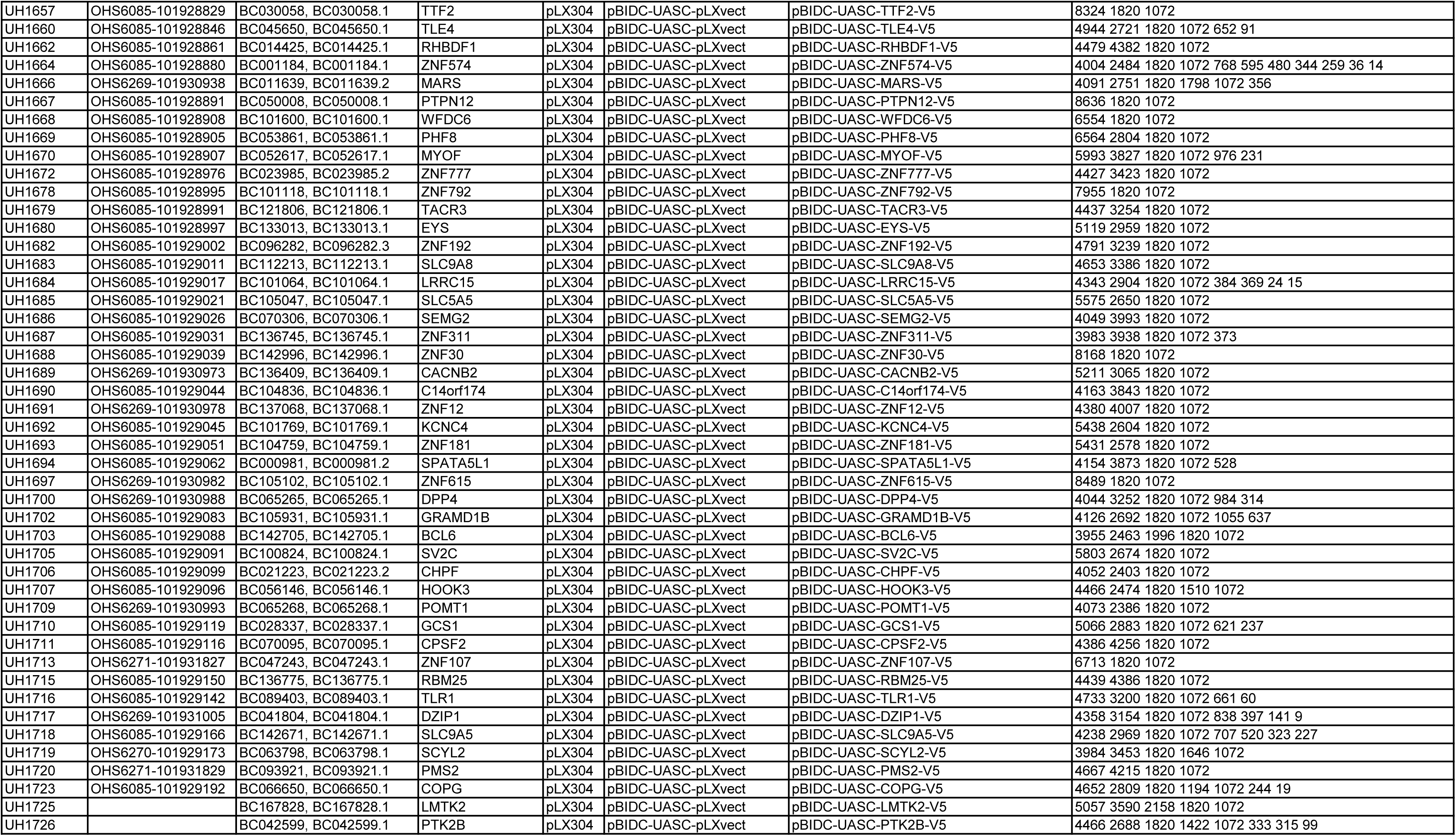
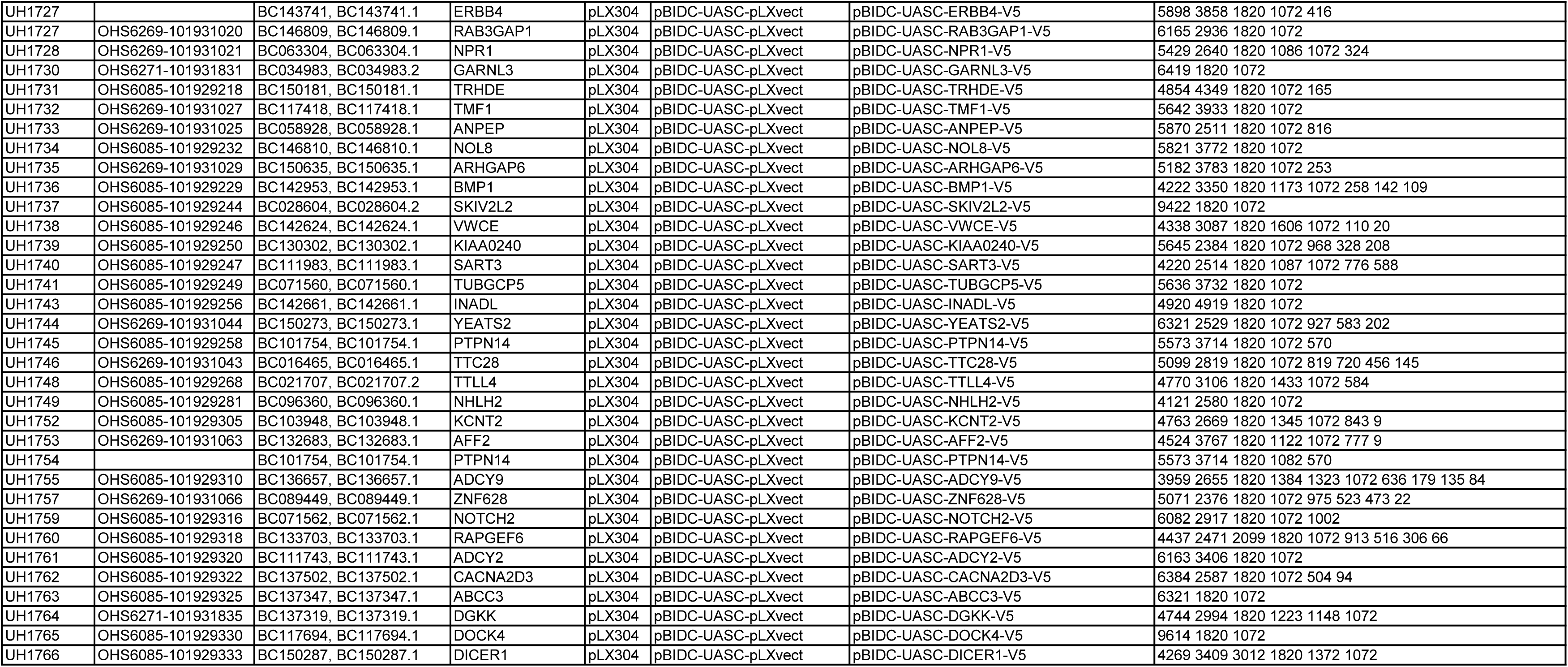
1,538 UAS-ORF constructs generated by CRISPRshuttle from 1,588 ORF clones of the human CCSB-Broad Lentiviral Expression Library.

